# Identification and characterization of the WYL BrxR protein and its gene as separable regulatory elements of a BREX phage restriction system

**DOI:** 10.1101/2021.12.19.473356

**Authors:** Yvette A. Luyten, Deanna E. Hausman, Juliana C. Young, Lindsey A. Doyle, Kerilyn M. Higashi, Natalia C. Ubilla-Rodriquez, Abigail R. Lambert, Corina S. Arroyo, Kevin J. Forsberg, Richard D. Morgan, Barry L. Stoddard, Brett K. Kaiser

## Abstract

Bacteriophage exclusion (‘BREX’) phage restriction systems are found in a wide range of bacteria. Various BREX systems encode unique combinations of proteins that usually include a site-specific methyltransferase; none appear to contain a nuclease. Here we describe the identification and characterization of a Type I BREX system from *Acinetobacter* and the effect of deleting each BREX ORF on growth, methylation, and restriction. We identified a previously uncharacterized gene in the BREX operon that is dispensable for methylation but involved in restriction. Biochemical and crystallographic analyses of this factor, which we term BrxR (‘BREX Regulator’), demonstrate that it forms a homodimer and specifically binds a DNA target site upstream of its transcription start site. Deletion of the BrxR gene causes cell toxicity, reduces restriction, and significantly increases the expression of BrxC. In contrast, the introduction of a premature stop codon into the BrxR gene, or a point mutation blocking its DNA binding ability, has little effect on restriction, implying that the BrxR coding sequence and BrxR protein play independent functional roles. We speculate that the BrxR coding sequence is involved in *cis* regulation of anti-phage activity, while the BrxR protein plays an additional regulatory role, perhaps during horizontal transfer.

## INTRODUCTION

Bacteria and phage are locked in an ancient conflict that has driven bacteria to evolve numerous and varied anti-phage defense systems, including restriction-modification systems (RM), CRISPR-Cas, abortive infection mechanisms (abi) and toxin-antitoxin (TA) gene pairs (1–7). These defense systems operate either by discriminating self-genomes from non-self and degrading foreign DNA (RM and CRISPR-Cas) or by inducing dormancy or a cell suicide response (*abi* and TA systems) (2).

BREX is a largely uncharacterized defense system that restricts phage by a poorly understood mechanism (8, 9). First described in *S. coelicolor* in the 1980s as being associated with a “phage growth limited” (pgl) phenotype (10), the BREX family was subsequently found to comprise at least 6 subfamilies that are widely distributed in bacteria and archaea (present in ∼10% of genomes) (2, 8). The various subtypes typically contain four to eight genes drawn from at least 13 gene families. DNA modification is a central feature of BREX function; five of the known subfamilies encode a site-specific DNA methyltransferase (PglX), while a sixth family encodes a PAPS reductase gene that also modifies host DNA (8). The most abundant family (BREX type 1) is present in ∼55% of BREX-containing genomes and encodes several large ORFs (>75 kDa) with identifiable enzymatic motifs, including an ATPase domain (BrxC), an alkaline phosphatase domain (PglZ), and a AAA+ ATPase (BrxL) with homology to Lon protease and/or RadA. How these factors collaborate in a BREX restriction system is not understood.

Two Type 1 BREX systems (from *E. coli* and *B. cereus)* and one Type 2 BREX system (from *S. coelicolor;* previously referred to as ‘phage growth limited’ or ‘Pgl’) have been described in detail. The timing and mechanism of phage restriction appears to differ between these systems. Bacteria harboring Type 2 BREX systems are sensitive to an initial round of phage infection (and permit lysogeny) but are then resistant to phage emerging from that cycle (10, 11). In contrast, type 1 systems appear to halt phage production prior to the first round of infection at a step very early after phage absorption and prevent lysogenization (8, 9). Both systems utilize PglX-directed DNA methylation to distinguish self from non-self, but do not appear to restrict phage by degrading their genome, and no BREX ORFs appear to encode a nuclease (8, 9), although one recently described BREX system (from *Escherichi fergusonii)* has been found to include a type IV restriction enzyme that appears to act independently of the function and mechanism of the surrounding BREX genes (12). BREX also does not appear to function by an abortive infection (abi) or cell suicide (TA) mechanism (8). Thus, the mechanism of BREX-mediated restriction appears to be novel from previously described defense systems.

BREX systems may employ various strategies to regulate or suppress inherently toxic activities during horizontal transfer and/or after establishment in an existing bacterial host until needed. Such fail-safe mechanisms have been described for other phage restriction systems. They include transcriptional repressors such as C proteins in R-M systems (which delay expression of a nuclease upon transfer to a new host) (13, 14), direct inhibitors of restriction factors such as the TRIP13 factor in CBASS restriction systems (which attenuate activity until detection of phage) (15), and phase variation mechanisms (16). In Type 2 BREX systems, PglX and PglZ appear to form a toxin/anti-toxin pair (17), and additional interactions between BREX factors have been proposed to regulate toxic activities (9). The Type 1 BREX system from *B. cereus* contains two operons (BrxA, BrxB, BrxC and PlgX on one; PglZ and BrxL on the other), further suggesting that BREX systems might be appropriately organized for transcriptional regulation (8). Finally, phase variation of PglX has been observed in the type II BREX system from *Streptomyces coelicolor* (17–19); a high frequency of disruptions in PglX coding sequences may indicate that phase variation may be a frequent mechanism of BREX regulation (8).

Two recent studies of host-virus dynamics in *Vibrio* and *Proteus* bacterial species, performed in India and Ireland, respectively, have further illustrated the global distribution and likely role of a broad family of related transcriptional regulators in various phage restriction systems including BREX (20, 21). Clinical isolates of *Vibrio cholera* contain a ∼100 kb mobile genetic element (the SXT integrative conjugative element) that encodes various defense systems (including BREX) within a genetic hotspot (20). Analysis of neighboring genes in 76 distinct isolates revealed that an ORF encoding a WYL domain is present upstream of the defense systems (20).

WYL proteins have also been previously linked to numerous CRISPR/Cas genetic loci (22) and shown to transcriptionally regulate CRISPR transcripts in several cyanobacterial species (23). WYL proteins also transcriptionally regulate numerous cellular pathways, including DNA damage response (24, 25). WYL domains are frequently encoded on the same polypeptide with two other domains—a DNA binding winged helix-turn-helix (wHTH) domain and a WCX domain-- and have been proposed to function as sensors that bind to environmental ligands -- perhaps phage-derived modified nucleotides or bacterial second messengers -- that allosterically activate the associated DNA binding domain on the same polypeptide (22–25). Currently, studies of multiple systems that contain and depend upon WYL protein domains have yet to identify an effector that clearly binds to and regulates their function.

In this study we characterize a bacterial Type 1 BREX system from *Acinetobacter,* a bacterial genus that includes the pathogenic *Acinetobacter baumanni* (including a recently described pathogenic, patient-derived strain that also contains a BREX system) (26). We identify its PglX methyltransferase target site, determine the requirements of BREX ORFs for PglX-directed methylation, and then further study the requirements of the same ORFs for phage restriction. We identify a 5’ ORF often linked to Type I BREX systems and show that this gene—which we call BrxR—is required for phage restriction but dispensable for methylation. We demonstrate that BrxR is a sequence-specific DNA binding protein comprising wHTH, WYL and WCX domains, propose that the protein and its reading frame act respectively as *trans* and *cis* transcriptional regulators of the BREX system, and suggest that its function has been widely adopted in more divergent bacterial species to regulate defense systems.

## MATERIAL & METHODS

### Identification and subcloning of a novel type I BREX system

Whole genome sequencing of *Acinetobacter* sp. NEB394 was carried out using Pacific Biosciences (PacBio) SMRT sequencing technology (27). The assembled genome and associated plasmids have been submitted to NCBI (NCBI:txid2743575). Putative DNA modification motifs were identified using PacBio SMRTAnalysis 2.3.0 Modification_and_Motif_Analysis_1.0. To assign methylation function to DNA methyltransferase genes, putative DNA methyltransferase genes identified within REBASE (28) and flanking genes were examined using the standard protein blast (BLASTP) program from NCBI (29). One putative N6A DNA methyltransferase appeared as part of a classic BREX system with flanking genes closely related to the BrxA, BrxB, BrxC, PglZ, and BrxL genes (8).

Oligonucleotide primers for cloning and mutagenesis and DNA constructs for binding assays (Supplementary Table S1) were synthesized by Integrated DNA Technology (Coralville, IA). Molecular biology reagents including Q5 Hot Start High-Fidelity DNA polymerase, NEBuilder HiFi DNA assembly master mix, restriction enzymes, DNA size standards, and competent cells were provided by New England Biolabs (NEB). Genomic DNA isolation, plasmid purification, agarose gel and PCR clean ups were performed using Monarch DNA kits (NEB). Plasmid DNA constructs were confirmed by sequencing on the ABI 3130xl capillary machine (Applied Biosystems) and PacBio RSII (Pacific Biosciences).

The 14.1kb BREX operon was PCR amplified as three ∼4.7 kB fragments from genomic DNA using overlapping primers designed for NEBuilder HiFi assembly. The expression plasmid pACYC184 (30) was amplified by inverse PCR to insert the BREX operon downstream of the Tet promoter. The *Acinetobacter* sp. NEB394 BREX system was initially subcloned in the pACYC184 expression vector under the control of a constitutive Tet promoter, using NEBuilder HiFi DNA assembly mix. Briefly, PCR-amplified insert and plasmid DNAs were mixed in equimolar ratios, incubated at 50°C for 60 minutes, and transformed into ER2683 (F’proA+B+ lacIq ΔlacZM15 miniTn10 (KanR) fhuA2 Δ(lacI-lacA)200 glnV44 e14-rfbD1 relA1 endA1 spoT1 thi-1 Δ(mcrC-mrr)114::IS10) competent cells per manufacturer’s instructions. Individual colonies were selected and grown overnight in LB broth (10 g/L soy peptone, 5 g/L yeast extract, 5 g/L NaCl, 1 g/L MgCl_2_, 1 g/L dextrose) supplemented with chloramphenicol (25 μg/ml). Total DNA was isolated from overnight cultures and prepared for sequencing on PacBio RSII (Pacific BioSciences, Menlo Park, CA). The BrxR-Stop and BrxR-R74A point mutation inserts were generated by assembly PCR with Accuprime Pfx polymerase (Invitrogen) using two large fragments amplified from the wildtype operon vector and two overlapping adapter oligos containing the desired mutations. The mutated inserts were cloned into the wildtype operon vector digested with BstZ17I (NEB) using NEBuilder HiFi assembly and the full plasmids were sequence-verified.

In a second round of subcloning to further investigate possible regulatory interactions between the BrxR protein and a putative *cis* regulatory sequence upstream of the *Acinetobacter* BREX transcript, the constitutive tet promoter was replaced by the insertion of a 97 basepair sequence that immediately precedes the BREX operon. The introduction of that sequence was carried out in two sequential rounds of PCR. First round amplifications were performed using Q5 Hot Start High-Fidelity DNA polymerase (NEB) and primer pairs brxRBD_for1/brxRBD_rev1 (wild-type BREX, BrxR-Stop, and BrxR-R47A) or brxRBD_delR_for1/brxRBD_rev1 (BrxR gene deletion) (Supplementary Table S1) under the following conditions: 25 cycles at T_a_ = 66°C and 9-minute extension. Primer pairs were designed to introduce 70 of 93 basepair sequence into the wild-type and point mutation BREX constructs and 61 of 93 basepair sequence into the BrxR gene deletion construct in the first round of amplification. Following amplification, template DNA was removed by digestion with DpnI for 30 minutes at 37°C. PCR products were analyzed by agarose gel electrophoresis and purified using Monarch PCR purification kit. Purified products were used as template in the second round of amplification. Primer pairs, brxRBD_for2/brxRBD_rev2 and brxRBD_delR_for2/brxRBD_delR_rev2 were designed to insert the remainder of the desired sequence upstream of the BREX operon (23 and 32 basepairs, respectively) and to contain 24 and 32 base overlaps for Gibson assembly post-amplification. Amplification was performed using Q5 Hot Start High-Fidelity DNA polymerase (NEB) under the same cycling parameters described for round 1 amplification. Following amplification, the desired ∼17 kB band was gel purified using Monarch Gel Purification kit and purified amplicons were self-ligated using NEBuilder HiFi DNA Assembly Master Mix and transformed into ER2683 competent cells.

Individual colonies were grown overnight in LB broth supplemented with chloramphenicol (25 μg/ml). Plasmid DNAs were isolated from overnight cultures for analysis by restriction digestion and PacBio Sequel II sequencing of the entire plasmid. The size of plasmid constructs was determined by digestion with HpaI in a 30 µl reaction containing 5 ng template DNA, 1x NEB CutSmart Buffer, and 1 µl restriction enzyme, at 37°C for 30 minutes. Digestion products were analyzed by agarose gel electrophoresis. Insertion of the 93 basepair sequence was verified via Sanger sequencing (ABI 3130xl capillary machine) and the entire construct sequence (∼17 kb) was confirmed via sequencing on the PacBio Sequel II system (Pacific Biosciences).

### Pacific Biosciences Sequencing

*In vivo* modification activities and sequence specificity of the wild-type BREX construct and subsequent gene deletion or point mutation plasmids were analyzed by sequencing on the PacBio RSII (Pacific Biosciences). Determination of *in vivo* DNA modifications and corresponding target motifs was performed using PacBio SMRTAnalysis 2.3.0 Modification_and_Motif_Analysis_1.0. Prior to library preparation, input DNA was sheared to an average size of 5-10 kb using gTubes (Covaris) and concentrated using 0.6V Ampure beads (Pacific Biosciences). Libraries were prepared according to manufacturer’s protocol using the SMRT-bell Template Prep kit 1.0 (Pacific Biosciences) for sequencing on the PacBio RSII (Pacific Biosciences).

### Transcription start-site (TSS) analysis

Three independent cultures of *Acinetobacter* sp. NEB394 were grown at 30°C in LB media to late log phase (OD_600_ between 0.7-0.75). Two volumes RNA protect (Qiagen) was added to the culture prior to RNA extraction using Monarch total RNA miniprep kit (NEB). Isolated RNA was quantified using Quibit RNA BR assay kit (Invitrogen) and used for Cappable-seq and SMRT-Cappable-seq.

Using the Cappable-seq method (31), two biological replicate enriched transcriptional start-site (TSS) libraries were generated. Briefly, a capping reaction was performed using vaccinia capping enzyme (NEB) to add a desthio-biotinylated GTP (DTB-GTP) cap to the 5’ triphosphate terminus of RNA. The capped RNA was fragmented using T4 polynucleotide kinase and primary RNA transcript isolated through 2 rounds of streptavidin magnetic bead enrichment. Following enrichment, the DTB-GTP cap was removed leaving a 5’ monophosphate terminus using RNA 5’pyrophophohydrolase (NEB), RNA was bound to AMPure beads and eluted in low TE (10mM Tris pH8.0, 0.1mM EDTA pH8.0). Eluted RNA was used for library preparation using the NEBNext small RNA library kit (NEB). The enriched and control libraries were amplified through 16 cycles and 10 cycles of PCR, respectively. RNA sequencing was performed on Illumina MiSeq*®* with single reads of 100 bases using V3 Illumina platform.

Single-end Illumina reads were trimmed to remove adaptors using cutadapt (32) with default parameters. Reads were mapped to the *Acinetobacter* sp NEB 394 plasmid pBspH3 (CP055285.1) using Bowtie2 v2.3.4.3(33). Gene annotations were derived from the NCBI *Acinetobacter* sp 394 plasmid pBspH3 annotation (GenBank: CP055285.1). Mapped reads were visualized using Integrative Genomics Viewer (34).

Total RNA from a third biological replicate was used to prepare a SMRT-cappable-seq library with minor modifications to the previously described method (35). Second-strand cDNA was bulk amplified with LongAmp HotStart Taq using CapSeq_for_dU_RM1 and CapSeq_rev_dU_RM1 primers. cDNA was size selected using Sage BluePippin (Sage Science) at a cut-off threshold of 2.5kb. Size-selected cDNA was PCR amplified, purified and PacBio SMRTbell adapters were ligated as previously described. Non-size-selected and size-selected SMRTbell libraries were sequenced using the PacBio RSII and Sequel platforms.

### Gene Deletions and point mutations

Individual genes were systematically removed from the cloned BREX operon (pAcBREX_WT) by inverse PCR. The pAcBREX_WT was amplified using Q5 Hot Start High-Fidelity DNA polymerase (NEB) and primers containing a 15-21 base overlap (see Supplementary Table S1) targeting DNA sequences immediately adjacent to the 5’ and 3’ ends of the gene to be removed. Following amplification, template DNA was removed by digestion with DpnI for 30 minutes at 37°C. PCR products were analyzed by agarose gel electrophoresis and purified using Monarch gel extraction or PCR purification kits. Purified amplicons were self-ligated using NEBuilder HiFi DNA Assembly Master Mix and transformed into ER2683 competent cells to generate a series of 7 plasmid constructs each lacking a single BREX gene (pAcBREX_ΔBrxR, pAcBREX_ΔBrxA, pAcBREX_ΔBrxB, pAcBREX_ΔBrxC, pAcBREX_ΔPglX, pAcBREX_ΔPglZ, pAcBREX_ΔBrxL).

Individual colonies were grown overnight in LB broth supplemented with chloramphenicol (25 μg/ml). Plasmid and total DNAs were isolated from overnight cultures for analysis by restriction digestion and PacBioRSII sequencing respectively. Restriction enzymes selected for screening contained 2 recognition sites within the pAcBREX_WT plasmid, 1 within and 1 outside of the gene to be removed. Plasmid constructs were digested with AleI (pAcBREX_ΔBrxR, pAcBREX_ΔPglZ), MluI (pAcBREX_ΔBrxA, pAcBREX_ΔBrxC), ApaLI (pAcBREX_ΔBrxB), KpnI (pAcBREX_ΔBrxL), and ZraI (pAcBREX_ΔPglX) in a 30 µl reaction containing 0.5 ng template DNA, 1x NEB CutSmart Buffer, and 1 µl restriction enzyme, at 37°C for 30 minutes. Digestion products were analyzed by agarose gel electrophoresis. The ligation junction for plasmids producing the predicted restriction patterns were confirmed via Sanger sequencing (ABI 3130xl capillary machine). Upon confirmation by restriction digestion and Sanger sequencing, total DNA derived from the corresponding overnight culture was used to generate PacBio libraries analyzed for *in vivo* modification activity and sequence confirmation of the expression plasmid.

### Bacterial growth and phage restriction assays

All infections using phage λ were performed with a virulent mutant of the phage unable to undergo lysogeny (λ_vir_) (36). In all experiments, cells were transformed with pACYC-based plasmids encoding the wild-type BREX operon, various deletion mutants, or control vectors that lack the BREX operon. Transformants were stored at -80^°^C as glycerol stocks. Experiments were performed both in *E. coli* 5-alpha cells (fhuA2 a(argF-lacZ)U169 phoA glnV44 a80a(lacZ)M15 gyrA96 recA1 relA1 endA1 thi-1 hsdR17; NEB) and in *E. coli* strain ER2683 ((F’proA+B+ lacIq ΔlacZM15 miniTn10 (KanR) fhuA2 Δ(lacI-lacA)200 glnV44 e14-rfbD1 relA1 endA1 spoT1 thi-1 Δ(mcrC-mrr)114::IS10; NEB).

All experiments began using isolated colonies derived from freezer stocks, which were then used to inoculate 4 ml of lysogeny broth (LB; 10 g/L casein peptone, 10 g/L NaCl, 5 g/L ultra-filtered yeast powder) supplemented with 1.25 mM MgCl_2_, 1.25 mM CaCl_2_, and 25 mg/ml chloramphenicol (‘overnight culture media’). Cultures were grown overnight at 37^°^C with shaking at 220 rpm. The next day, overnight cultures were diluted 50-fold in 4 ml of LB supplemented with 1.25 mM MgCl_2_, 1.25 mM CaCl_2_, 0.2% maltose, and 25 µg/ml chloramphenicol (‘outgrowth media’) and grown to mid-log (absorbance readings at 600 nm varied between 0.2 and 0.6) at 37^°^C, shaking at 220 rpm. Mid-log cultures were then diluted to an OD_600_ of 0.01 in LB containing 1.25 mM MgCl_2_, 1.25 mM CaCl_2_, 25 mg/ml chloramphenicol. Phage titers were determined via a plaque assay to ensure accurate multiplicities of infection (’MOIs’). Phage λ_vir_ samples were serially diluted in SM buffer (50 mM Tris-HCl, 25 mM NaCl, 4 mM MgSO_4_) before use and added to the bacterial samples at MOIs ranging from 0.001 to 1.0. SM buffer alone was added to samples without phage. Cultures were then arrayed in triplicate across a 96-well plate (Greiner cat#655083, 100 µl per well) and grown for ten hours in a BioTek Cytation three plate reader at 37°C with continuous orbital shaking at 282 cycles per minute. Absorbance readings were taken every 15 minutes at 600 nm.

Additional, complementary phage plaque formation assays were performed using the same titred phage stock and overnight cultures of ER2683 grown in overnight culture media. Multiple dilutions of the overnight cultures were generated the next morning (between 30- and 100-fold) in outgrowth media and then grown to an OD_600_ of approximately 0.4 to 0.5. 80 microliters of each sample were mixed with 3 mL of top agar (0.5% agar in LB supplemented with 1.25 mM MgCl_2_, 1.25 mM CaCl_2_, and 25 µg/ml chloramphenicol) and applied to bottom agar plates (1.5% agar in LB supplemented with 1.25 mM MgCl_2_, 1.25 mM CaCl_2_, and 25 µg/ml chloramphenicol). Plates were allowed to solidify and dry for approximately 15 minutes and then spotted with 5 µl of 10-fold serial dilutions (ranging from 10^1^ to 10^8^ dilutions) of titred phage. Plates were incubated overnight at 37°C and examined the following morning for plaque formation. Each experiment was performed with a minimum of three biological replicates.

### Protein expression and purification

The BrxR gene was subcloned from the native Acinetobacter 394 locus into the pET15b plasmid to contain an N-terminal, thrombin-cleavable 6XHistidine (His6) tag. Inductions were carried out in BL21 (DE3) pLysS *E*. *coli* cells. For inductions, a 10-mL overnight starter culture was grown in LB media with ampicillin (100 μg/mL), diluted 100-fold into the same media and incubated at 37°C with shaking until an OD_600_ of 0.6 was reached. IPTG was added to a final concentration of 500 µM, and the culture was shaken for an additional 18-22 hours at 16°C. Cell pellets were harvested by centrifugation and stored at −20°C.

Cell pellets were resuspended in lysis buffer (25 mM Tris (pH 7.5), 300 mM NaCl, 20 mM imidazole)), lysed by sonication on ice and centrifuged in an SS34 rotor for 25 minutes at 18,000 rpm. The supernatant was filtered through a 5 μm syringe filter, and the clarified lysate was incubated in batch with Ni-NTA agarose resin (Qiagen) for 1 hour at 4°C with rotation. The resin was transferred to a gravity filtration column (Bio-Rad), washed with >50 volumes of lysis buffer at 4°C and eluted in lysis buffer supplemented with 200 mM imidazole. Biotinylated thrombin (EMD Millipore) was added to the eluted protein (1 unit of thrombin per mg of protein), and the sample was dialyzed into 200 mM NaCl, 50 mM Tris (pH 7.5) overnight at 4°C. Removal of the His6 tag was assessed by running “pre-thrombin” and “post-thrombin” samples on a 4-12% BOLT SDS PAGE gel in MES buffer (Invitrogen); the difference in size between the His6-tagged and untagged version was clearly resolved on these gels. The sample was concentrated to 2 mL in an Amicon Ultra centrifugal filter (10,000 molecular weight cutoff; Millipore). Thrombin was removed by incubation with streptavidin agarose (Novagen) for 30 minutes at 4°C. The sample was filtered through a 0.22 μm centrifugal filter and loaded onto a HiLoad 16/60 Superdex 200 prep grade size exclusion column (Millipore Sigma) equilibrated in 25 mM Tris (pH 7.5), 200 mM NaCl. Peak fractions were pooled and concentrated to 14 mg/mL. Single-use aliquots were flash frozen in liquid nitrogen and stored long-term at -80°C. This untagged BrxR (i.e with the His6 tag removed) was used in biochemical and crystallography experiments.

To produce selenomethionyl (SeMet)-containing BrxR for crystallization, we followed a previously described induction and growth protocol (37) using the same plasmid, *E. coli* strain and purification strategy described above. This protein was concentrated and stored at 12 mg/mL.

### DNA binding assays

DNA substrates for gel shift assays were amplified by PCR using Q5 Polymerase (NEB) and purified using a DNA Clean and Concentrator kit (Zymo Research). DNA was used at a final concentration of 20 nM in binding experiments. Untagged BrxR proteins were diluted in 100 mM NaCl, 25 mM Tris (pH 7.5) and used at final concentrations ranging from 25 nM to 800 nM. Binding reactions were assembled in binding buffer (20 mM Tris (pH 7.9), 40 mM NaCl, 2.5% sucrose)) and incubated at 22°C for 30 minutes. Samples were resolved on native acrylamide gels containing 0.5X TBE buffer and 7% acrylamide (made from a stock of 40% acrylamide with a ratio of 29:1 acrylamide:bisacrylamide; Bio-Rad). The running buffer for the gels was 1XTBE. Prior to loading samples, the gels were pre-run for 30 minutes at 90 volts in 1 X TBE buffer and the wells were flushed using a syringe. 4 µl of each binding reaction was loaded on the gel (without dye) and run at room temperature for 100 minutes at 90 volts. Gels were stained for 30 minutes in SybrGold (Invitrogen) diluted in 1X TBE and visualized on a Typhoon scanner using a 488 nm laser and Cy2 filter.

### BrxR crystallographic structure determination

All crystallographic data and refinement statistics are provided in Table 1. The structure of apo BrxR was initially determined to 2.3 Å resolution via single anomalous dispersion (SAD) phasing using SeMet-derivatized protein. The crystals were grown by hanging drop vapor diffusion in drops set with 1 µl of protein (12 mg/mL) plus 1 µl of well solution (10% PEG3000, 0.1 M HEPES (pH 7.3), 2% benzamidine-HCL). For cryopreservation, crystals were transferred into crystallization solution containing 20% ethylene glycol and 0.1% H_2_O_2_, incubated for one minute and flash frozen in liquid nitrogen. Diffraction data for the SeMet-containing BrxR was collected at the Advanced Light Source synchrotron facility (ALS, Berkeley, CA) at beamline 5.0.2. A single-wavelength data set was collected with an incident x-ray wavelength of 0.9792 Å, corresponding to the anomalous dispersion peak for selenium (12.662 keV).

**Table 1.** Crystallographic data and refinement statistics.

| PDB ID | BrxR (unbound)<br>7T8L | BrxR (DNA bound)<br>7T8K |
| --- | --- | --- |
| **Data Collection** |  |  |
| Space group | P 1 21 1 | C 1 2 1 |
| Unit cell |  |  |
| a, b, c | 65.3, 48.67, 96.11 | 151.6, 71.5, 87.4 |
| alpha, beta, gamma | 90, 102.28, 90 | 90, 105.1, 90 |
| Wavelength (Å) | 0.9795 | 0.9763 |
| Resolution range (Å) | 93.7-2.08 (2.12-2.08) | 64.2-2.3 (2.38-2.3) |
| Unique reflections | 35573 | 38731 |
| R-merge | 0.066 (0.239) | 0.075 (0.572) |
| R-meas | 0.091 (0.319) | 0.105 (0.668) |
| R-pim | 0.051 (0.176) | 0.042 (0.264) |
| CC1/2 | 0.996 (0.862) | 0.975 (0.847) |
| I/sigma(I) | 12.27 (1.3) | 11.3 (2.6) |
| Chi^2 | 1.114 (0.779) | 1.375 (0.655) |
| Multiplicity | 3.0 (3.2) | 6.2 (6.1) |
| Completeness (%) | 99.7 (99.8) | 97.5 (100.0) |
| Mean B value | 30.902 | 48.65 |
| **Refinement** |  |  |
| Resolution (Å) | 63.92-2.0 | 64-2.3 |
| Completeness for resolution used (%) | 99.45 | 95.25 |
| R-work | 0.1910 | 0.1958 |
| R-free | 0.238 | 0.2376 |
| Number of non-hydrogen atoms | 4564 | 5588 |
| macromolecules | 4250 | 5445 |
| ligands | 156 | 26 |
| solvent | 158 | 117 |
| Protein residues | 504 | 563 |
| RMS(bonds) | 0.009 | 0.004 |
| RMS(angles) | 1.571 | 0.73 |
| Ramachandran favored (%) | 97.02 | 97.5 |
| Ramachandran allowed (%) | 2.58 | 2.5 |
| Ramachandran outliers (%) | 0.4 | 0 |
| Clashscore | 4.83 | 12.84 |
| Average B-factor | 34.65 | 55.04 |
| macromolecules | 33.42 | 55.04 |
| ligands | 35.35 | 68.08 |
| solvent | 35.19 | 51.91 |

Data were indexed and scaled using HKL2000 software (38). Experimental phases were determined by SAD phasing in CC4Pi’s CRANK phasing and building pipeline (39), producing an initial model with refinement Rwork and Rfree values of 0.24 and 0.28, respectively. Subsequent rounds of building were carried out in Coot (40) and refinement was performed using Refmac5 (41).

To visualize the DNA-bound form of BrxR, ssDNA oligonucleotides containing BrxR’s putative binding site (ATACCGTAAAAATAATTTACTGTAT top strand; bottom strand is complementary) were synthesized, annealed to produce blunt-end double strand DNA (dsDNA), and purified by HPLC (Integrated DNA Technologies). 150 µM dsDNA was incubated with 125 µM BrxR and crystallized by hanging drop diffusion in 0.04 M Citric acid, 20% w/v PEG 3350, 60 mM BIS-TRIS propane (pH 6.4). Crystals were cryopreserved in the same solution containing 22.2% ethylene glycol prior to freezing in liquid nitrogen. A native dataset was collected on ALS Beamline 5.01. The structure of the DNA-BrxR complex was determined by molecular replacement at 2.3 Å resolution using PHENIX (42), with the previously determined structure of the unbound protein as a molecular probe and phasing model. One copy of the protein homodimer was identified in the crystallographic asymmetric unit, with LLG and TFZ scores of 423.6 and 24.7, respectively. Density corresponding to the bound DNA target site was immediately identifiable in electron density difference maps, and the corresponding bound DNA was then manually fit using COOT (40). The structure of the DNA-bound BrxR complex was refined using PHENIX (42).

## RESULTS

### Identification and characterization of a novel type I BREX system and its PglX methyltransferase

We initiated this project to better understand the role of DNA methylation in *Acinetobacter* with the intent of identifying all methylated sites in a representative genome as well as the DNA methyltransferases (MTases) responsible for those modifications. For this purpose, we chose bacterial strain NEB 394 (*Acinetobacter species* NEB394), which was known to encode several R-M systems and carried out PacBio genome sequencing and corresponding methylome analysis of its genomic DNA. Identified among several uniformly methylated sites was a non-palindromic sequence corresponding to 5’-GTAG(A^m6^)T-3’, with methylation observed at the adenine base in the fifth position of the motif. Motif Software called 99.8% of the sites in the genome as methylated (1318 of 1320 sites) at 180X coverage, indicating essentially complete modification (Figure 1a).

**Figure 1.**
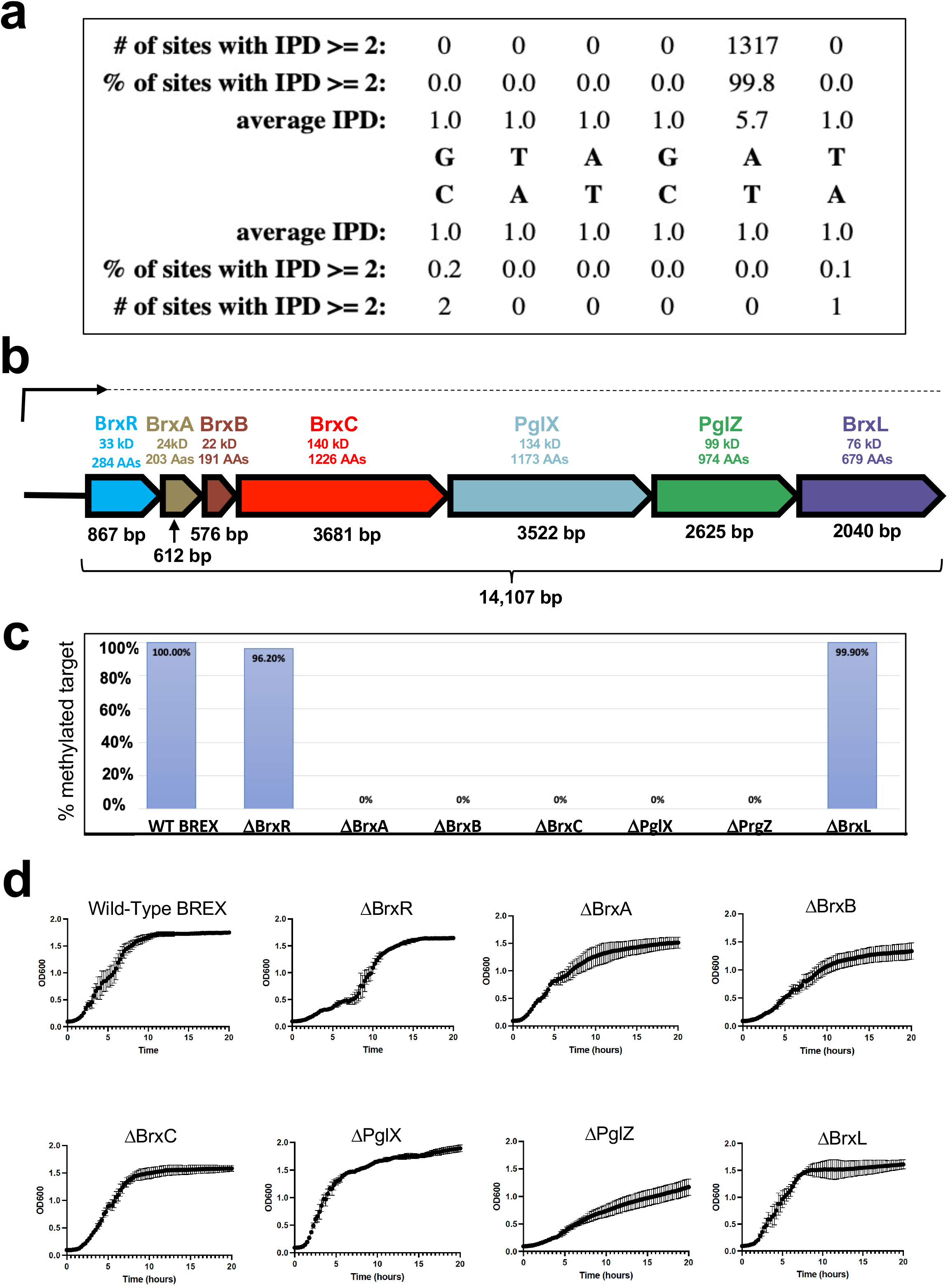
Type I BREX operon from Acinetobacter strain NEB394. ***Panel a:*** PacBio sequencing of genomic DNA from *Acinetobacter species* NEB394 harboring the endogenous BREX operon identifies the BREX target site and base (5’-GTAG**<u>A</u>**T-3’) methylated via action of the PglX methyltransferase. ***Panel b:*** Seven genes within the BREX locus encode for proteins ranging in size from 22 kD (BrxB, 191 residues) to 140 kD (BrxC, 1226 residues). A transcription start site (’TSS’) analysis shows that BREX is expressed via a single long transcript (upper arrow and dashed line) in logarithmically grown *Acinetobacter394* cells in the absence of a phage challenge. ***Panel c:*** Systematic deletion of each BREX gene from the pACYC-BREX plasmid and additional PacBio methylation analysis in transformed *E. coli* cells demonstrates that four BREX genes (BrxA, BrxB, BrxC and PglZ) are required for genome modification in addition to the PglX methyltransferase. Y-axis indicates the fraction of PglX target sites observed to be methylated in E. coli strains harboring each BREX genotype indicated on the X-axis. ***Panel d:*** Growth curves of NEB2683 *E. coli* cells transformed with pACYC-BREX plasmids in which each BREX gene has been systematically deleted. Three deletions (ΔPglZ, ΔBrxB and ΔBrxR) appear to display lags or reductions in growth rates. These observations are reflected in reduced transformation efficiencies and small or irregular colony size for the same constructs (Supplementary Figure S1).

Next, we compiled a list of DNA MTases present in the *Acinetobacter sp* NEB394 genome (encoded either on the chromosome or on plasmids), based on sequence similarity to known DNA MTases in the REBASE bioinformatic pipeline (28), that might target this motif. One plausible candidate was a gene appearing to encode a gamma-class amino DNA methyltransferase, members of which have been shown to hemi-methylate non-palindromic DNA target sites (43). To test the activity of this candidate, we subcloned the MTase gene and introduced it into *E. coli* under control of a constitutive promoter and then analyzed the methylome of the *E. coli* chromosome by PacBio sequencing. However, we observed no novel methylation events at the same sequence motif (or any others) via this analysis.

We then more carefully examined the genomic locus surrounding the candidate MTase to identify additional neighboring genes that might be required for the methylation activity. This analysis revealed that the MTase was flanked by genes known to be conserved across type I BREX phage restriction systems, indicating that it is likely a BREX-associated ‘PglX’ methyltransferase enzyme. The BREX system from *Acinetobacter* 394 contained the same conserved six genes present in all Type I systems, as well as a seventh gene located immediately upstream of the BREX locus (Figure 1b); we eventually termed this gene BrxR (‘BREX regulator’).

To determine if this BREX gene cluster (including PglX) was sufficient to methylate the target motif, we cloned all seven genes from *Acinetobacter* genomic DNA into the pACYC184 vector downstream of a constitutive tet promoter and introduced the plasmid into *E. coli* strain ER2683. PacBio sequencing and methylome analysis of the resulting transformed *E. coli* indicated full methylation at the GTAG**<u>A</u>**T(D) motif previously observed in the native *Acinetobacter sp* NEB394 host strain, with the Motif software calling 100% (1024 of 1024) of the GTAGAT sites in the *E. coli* genome (accession CP093221) as methylated at 116X coverage (Figure 1c). Note that BREX sites followed by a C overlap with, and are methylated by, *E. coli* dam MTase, GATC, and were thus excluded from the count of GTAGAT sites modified) PglX-mediated methylation has been reported for three bacterial species (*E. coli, B. cereus and L. casei)* (8,9,44); all recognize 6 basepair, non-palindromic motifs and methylate adenine in the fifth position of those targets, as we observe for *Acinetobacter 394* PglX.

### Four additional BREX genes are required for host methylation in E. coli by PglX

The observation that *Acinetobacter* 394 PglX was not functional when expressed by itself in *E. coli* suggested that it requires other proteins in the BREX system to form a functional DNA MTase. To investigate further, we generated seven additional pACYC vectors, each harboring the complete BREX system with one individual gene precisely deleted, transformed each of them into *E. coli* and performed additional PacBio sequencing and methylome analysis on the host *E. coli* genomic DNA (Figure 1c). Deletion of BrxR or BrxL (the last ORF in the BREX gene cluster) did not affect methylation at the target motif. In contrast, deletion of any one of the individual BrxA, BrxB, BrxC or PglZ genes resulted in loss of target methylation, implying that these proteins are all required, along with PglX, to form a functional DNA MTase. These findings are intriguing, as PglX by itself appears to have all the required domains for methyltransferase function and in most RM systems would be expected to be fully functional. The requirements of other BREX genes for PglX-mediated methylation has only been reported for *E. coli* (9). In that system, which comprises six genes (and does not include BrxR), all BREX components except BrxA and BrxL are required for methylation. The disparity in the requirement of BrxA between *Acinetobacter sp* NEB394 and *E. coli* may reflect differences in mechanism or regulation between the two systems. In all BREX systems it appears that host-protective methylation requires a multi-protein complex consisting of BREX factors. Such a requirement has also recently been described in the ciliate *Oxytricha*, where where multiple polypeptides tetramerize to form an active DNA methyltransferase.(45).

### Deletion of BrxR, BrxB or PglZ within the BREX operon delays or impairs cell growth

Further growth analyses of *E. coli* transformed with the same pACYC constructs described above (intact BREX or BREX harboring individual gene deletions) indicated that deletion of BrxR, BrxB and PglZ resulted in slower cell growth and/or an extended lag phase before approaching saturation (Figure 1d). The difference between those three BREX gene deletions was also visible when examining the size distribution of individual colonies arising on transformation plates (Supplementary Figure S1). Toxicity caused by deletion of PglZ has also been reported in the type II BREX system from *S. coelicolor,* in which PglZ is proposed to function as PglZ as an antitoxin partner to PglX (17).

### The BREX system restricts a lytic phage in E. coli when preceded either by a constitutive tet promoter or by the native upstream genomic sequence from Acinetobacter

An analysis of the BREX system in *Acinetobacter* 394 indicated that it was transcribed as a single mRNA, with its transcription start site (TSS) located 23 basepairs upstream of the BrxR gene (Figure 2a **and** Supplementary Figure S2). We therefore examined the ability of the BREX operon to restrict phage infection and lysis in *E. coli* when preceded either by a constitutive tet promoter or instead by the 97 bp sequence corresponding to the upstream *Acinetobacter* 394 genomic region immediately preceding BrxR (which included a predicted bacterial promoter immediately upstream of the *Acinetobacter* BREX TSS). For these experiments we conducted both plaque formation assays and bacterial growth and lysis assays in liquid culture, and employed a virulent mutant of *λ* phage, termed ‘λ_vir_’, which is unable to undergo lysogeny (36) against *E. coli* strain ER2683 (an MM294 derivative lacking all known methylation-active restriction systems (46)). The results indicated that in either genetic configuration, the BREX operon restricted phage infection and lysis by approximately equivalent amounts (Figure 2b,c).

**Figure 2.**
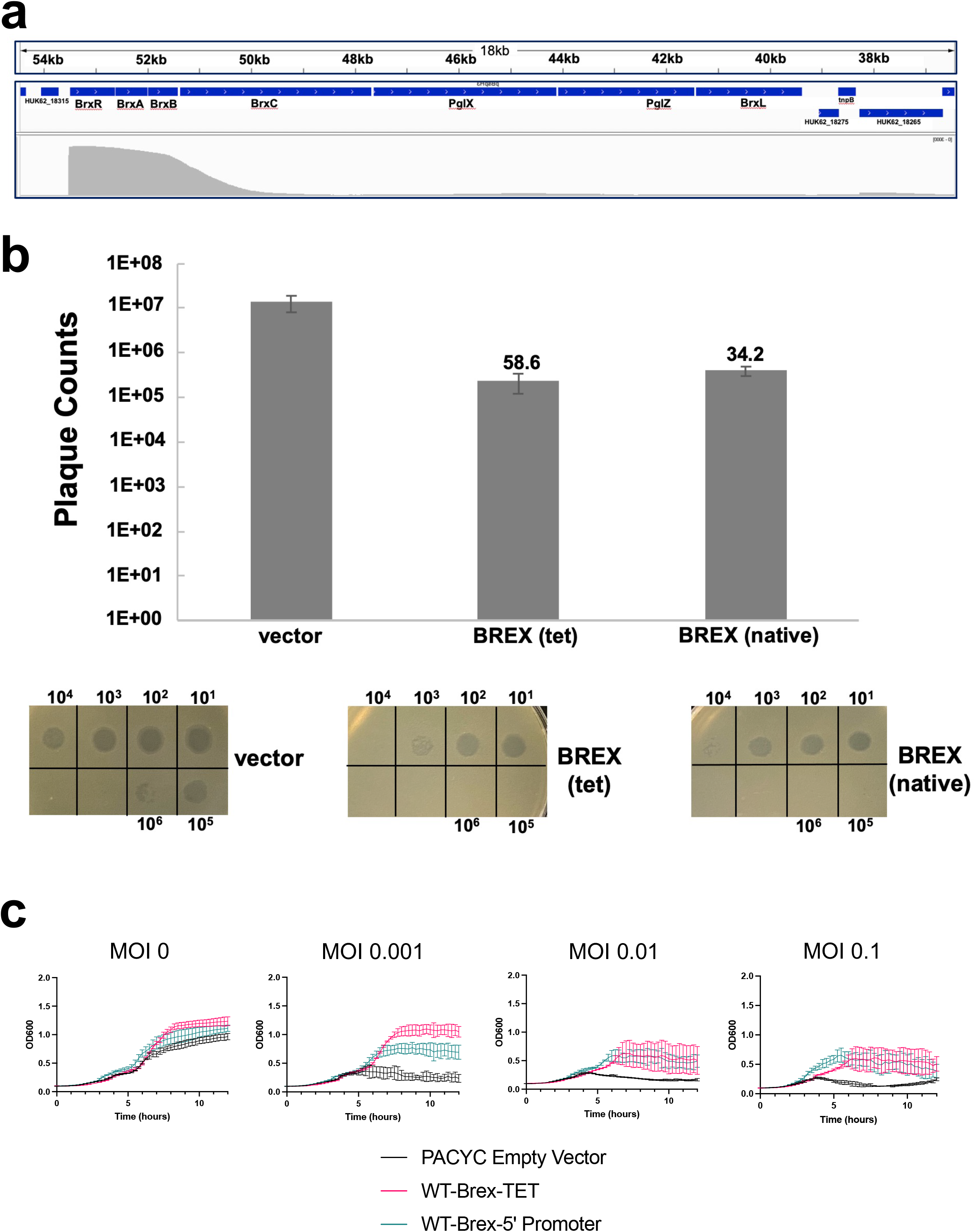
BREX is active in *E. coli* preceded either by a constitutive tet promoter or by the upstream genomic sequence from *Acinetobacter*. ***Panel a***: BREX transcription start site analysis in *Acinetobacter sp* NEB394. A single long transcript is detected beginning 23 base pairs upstream of the BrxR start codon. Numbers in kb show location within the pBspH3 plasmid containing the BREX system. The top track shows BREX ORFs. The bottom track shows PacBio Single Molecule Real time (’SMRT’) cappable sequence read coverage (gray, scale 0 to 3000). This shows a strong TSS just before BrxR and the BREX operon, with very low levels of internal operon reads, indicating one TSS for the entire operon. See Supplementary Figure S2 for additional detail. ***Panel b***: Restriction by BREX visualized in a plaque formation assay. Restriction was assayed using a plaque formation assay with λ_vir_ phage deployed against *E. coli* strain ER2683. For each construct, serial 10-fold dilutions of phage were spotted on a bacterial lawn, and individual plaques were counted. All assays were repeated in biological triplicate. The values above the bars indicate the fold-reduction in plaquing efficiency in the presence of the BREX operon preceded by a constitutive tet promoter (‘tet’) or by the upstream putative promoter and regulatory region from *Acinetobacter* (‘native’). Both operons generate similar (30- to 60-fold) reductions in plaquing efficiency, relative to the empty pACYC vector. ***Panel c***: Restriction by the same BREX constructs in liquid culture. Bacterial cultures were challenged with λ_vir_ phage at the indicated MOI and culture density (OD_600_ nm) was monitored over time.

To further examine the generalizability and reproducibility of these results, we tested the ability of the BREX operon to restrict phage infection and lysis in a different E. coli strain (NEB-5*α*). BREX again demonstrated a strong protective effect out to ten hours post-challenge, while cells lacking BREX displayed significant lysis and crash of the culture within five hours of the challenge. The level of protection conferred by the intact BREX system displayed a significant dependence on the MOI during the initial challenge; cells containing BREX eventually crashed at the highest MOI (0.1) but maintained continued growth at lower MOI’s of 0.01 and 0.001 (Supplementary Figure S3).

### All seven BREX subunits, including BrxR, are required for restriction of a lytic phage

We next went on to examine the effect of individually deleting each of the seven BREX genes from the operon in strain ER2683, using both plaque formation assays (Figure 3a) and liquid culture bacterial growth and lysis assays (Figure 3b and Supplementary Figure S4). Unlike the effect of each of the same deletions on host methylation (for which BrxR and BrxL were dispensable) deletion of each of the seven BREX genes led to a significant reduction in phage restriction, with deletion of each impairing phage restriction in liquid culture at both high and low phage MOIs (1.0 and 0.01).

**Figure 3.**
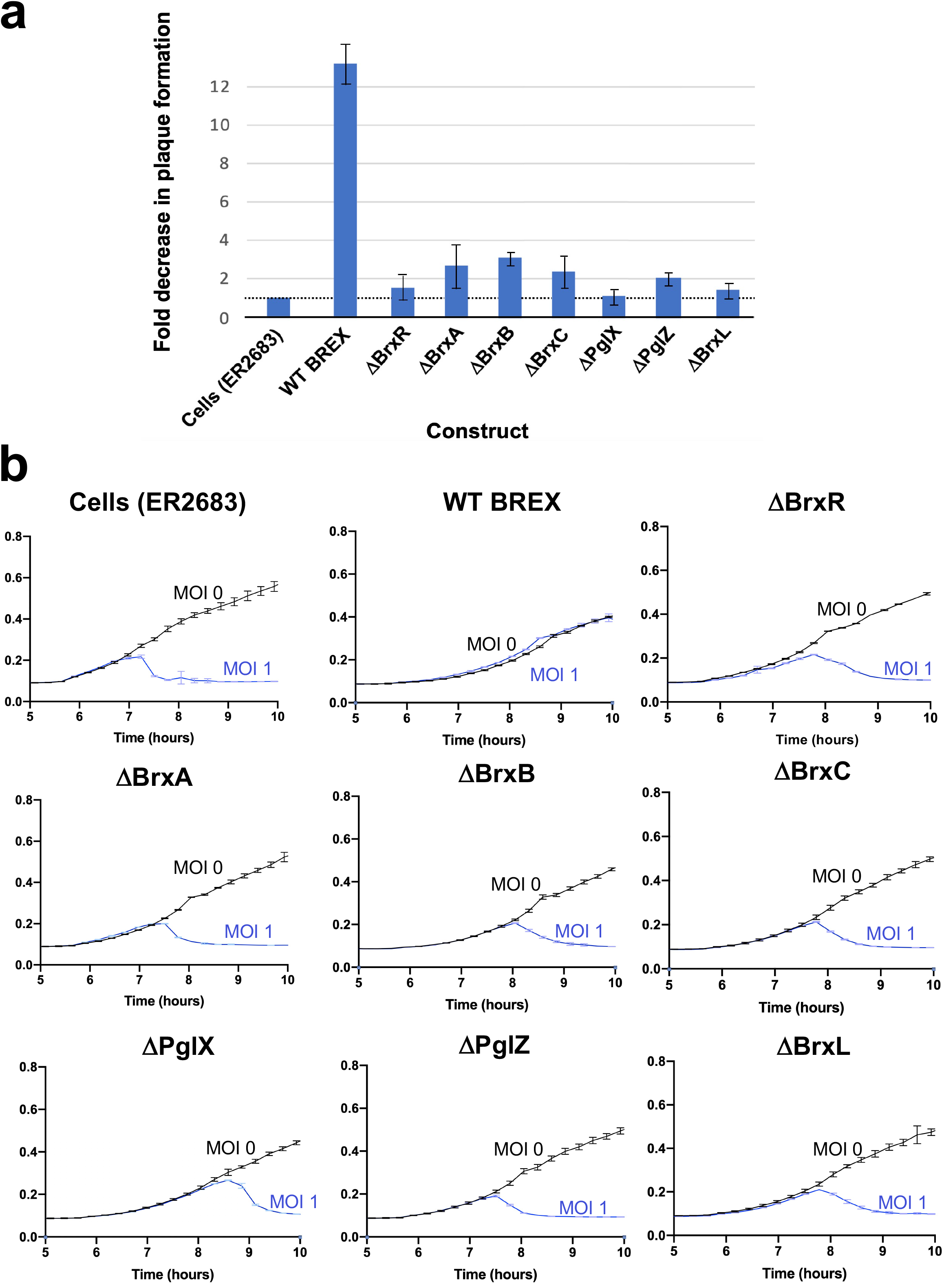
BREX restriction when each gene is precisely deleted. ***Panel a:*** Fold change in phage plaque formation using λ_vir_ phage and *E. coli* strain ER2683 as a function of the presence or absence of each gene in the BREX operon. Cells transformed with the indicated pACYC constructs, growing at log phase, were mixed with 0.5% top agar, plated on chloramphenicol plates to form a lawn, then spotted with 10-fold serial dilutions of λ_vir_ phage. The plaque formation efficiency of λ_vir_ phage on a ‘cells only’ (no BREX) control is normalized to 1; the corresponding fold reduction in plaquing formation efficiency of the same phage is then indicated for WT BREX and for BREX harboring a precise deletion of each gene. All seven genes appear to be involved in BREX restriction function. ***Panel b:*** Growth of *E. coli* strain ER2683 (New England Biolabs) challenged with the λ_vir_ phage at an MOI of 1.0. The intact BREX system conferred robust protection against λ_vir_ phage, whereas individual deletions of each BREX gene led to a significant reduction in phage restriction at both high MOI (shown here) and a lower MOI of approximately 0.01 (Supplementary Figure S4).

The effect of single gene deletions on phage restriction has also been reported for strain *E. coli* HS. In that system, all BREX genes except BrxA were required for phage restriction (9). Individual BREX gene requirements from other systems have not yet been reported.

### BrxR is a homodimeric winged helix-turn-helix-WYL-WCX protein with site-specific DNA binding function

We were intrigued by the observation that BrxR was required for BREX-mediated phage restriction, since it has not been reported as a core component of Type I BREX systems and is not part of the Type I systems so far described from *E. coli* or *B. cereus* (8, 9). Bioinformatic analyses indicated that BrxR contains three domains: an N-terminal winged helix-turn-helix (wHTH) domain, a WYL domain and a ‘WYL C-terminal extension’ (‘WCX’) domain. Proteins with this domain architecture are widely present in bacterial genomes and are thought in most cases to function as transcriptional regulators (25). The only structurally characterized HTH-WYL-WCX homolog, PafBC, is a transcriptional regulator of DNA damage responses in mycobacteria (25, 47). PafBC is notably different from most HTH-WYL-WCX proteins, including BrxR homologs, in that it contains a tandem duplication of HTH-WYL-WCX domains on the same polypeptide (proteins of this class represent only ∼5% of all HTH-WYL-WCX proteins).

We purified and crystallized BrxR as described in ‘Methods’ and determined its structure to 2.3 Å resolution (Figure 4 and Table 1). The protein forms a symmetrical homodimer comprising two intertwined protein chains, each of which contain an N-terminal wHTH domain, a central WYL domain, and C-terminal WCX domain (consistent with bioinformatic predictions) (Figure 4a,b and Supplementary Figure S5). The two exposed helices of the wHTH domain (a fold often associated with DNA binding) are positioned and spaced appropriately to interact with DNA at positions separated by approximately one full turn of a B-form double helix. The pair of beta strands that form the “wing” in this domain could also be positioned to contact DNA, as has been shown for other proteins of this class (48). The WYL domains are domain-swapped within the homodimeric structure of the protein, with each domain closely associated with the other subunit’s underlying wHTH domain. The core of the homodimer is further stabilized by interaction between helices in each protein subunit that are located between the wHTH and WYL domains (residues 71-119; gray in Figure 4a). The WCX domain, while somewhat diverged from that previously visualized for the PafBC transcriptional regulator (25), still forms the same overall fold. We therefore conclude that BrxR is a member of the HTH-WYL-WCX superfamily previously described in that prior analysis.

**Figure 4.**
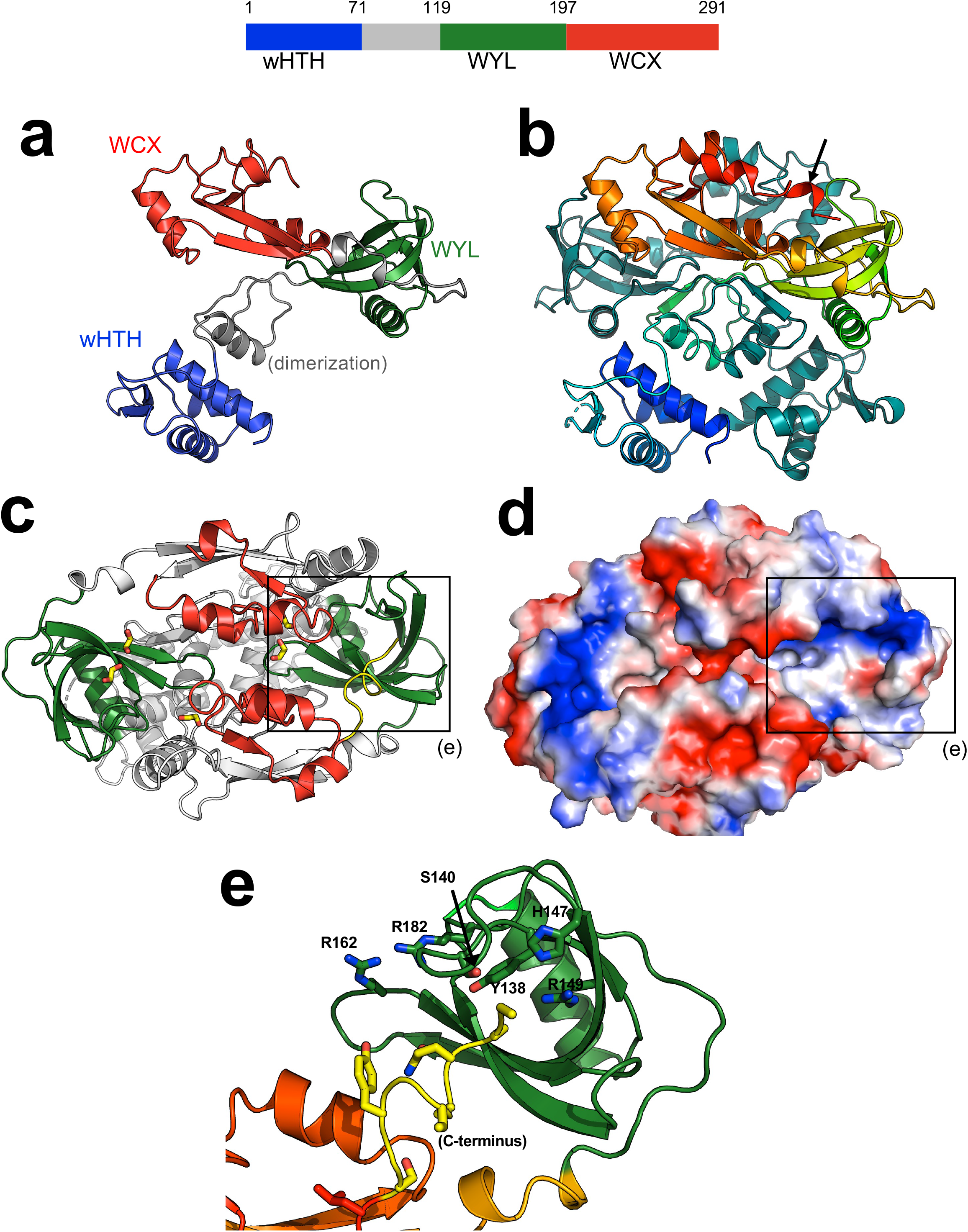
Crystal structure of BrxR. ***Top:*** Domain boundaries of BrxR. Residues 71-119 (gray) are involved in homodimer formation. See also Supplementary Figure S5. ***Panel a:*** Topology of one subunit of the BrxR dimer, colored as shown in the top schematic. ***Panel b:*** Structure of the BrxR homodimer, with one subunit colored as a spectrum ranging from the blue N-terminus to the red C-terminus and the second subunit colored in dark teal for contrast. The protein forms an intertwined, domain swapped homodimer. ***Panels c and d:*** Top-down views, respectively as a ribbon diagram and a charged electrostatic rendering, of the protein surface opposite the wHTH domains. The WYL domains (green in panel c) are at opposite ends of the protein and form a pair of positively charged pockets (blue color in panel d). Bound cryoprotectant ethylene glycol molecules observed in the interface between protein subunits and in the WYL pockets are indicated with yellow sticks (panel c). The boxes outline the WYL domain, and its pocket illustrated in panel e. ***Panel e:*** Close up view of the pocket formed by a WYL domain and multiple conserved residues in that region. Density for the C-terminus (yellow) is very weak, indicating high flexibility for that portion of the protein.

Inspection of the BrxR surface directly opposite its wHTH domains (Figure 4 c,d,e) revealed a pair of symmetry-related, basic pockets flanked in part by conserved residues from the WYL domain (including S140, Y138, H147, R149, R162, and R182). The pocket and its conserved residues corresponds to the same region that is predicted to mediate cofactor binding in the WYL domain of the PafBC transcription factor (25). The C-terminal end of one BrxR subunit (ending at amino acid 291) inserts into the pocket of the WYL domain of the opposite subunit (Figure 4e). The electron density for that C-terminal tail is weak (and entirely unobservable for the same C-terminal residues in the opposing subunit), indicating that this region is flexible and partially disordered, and could potentially be displaced by the binding of a (currently unidentified) effector molecule.

The structure of BrxR suggested that it might act as a DNA-binding protein, and potentially act as a transcriptional regulator. If so, given its structural symmetry it would likely bind a symmetric or nearly symmetric DNA target associated with the BREX operon. To test this possibility, we generated a panel of 13 DNA targets (mostly focused on upstream regions of BREX ORFs), each 250-300 bp in length, and tested the ability of purified BrxR to bind them in electrophoretic mobility shift assays (EMSAs or ‘gel shifts’). We observed low affinity interactions to all those DNA probes that appeared to correspond to non-specific interactions with DNA (Supplementary Figure S6).

Further examination of the *Acinetobacter* genomic sequence upstream of the BrxR ORF revealed the presence of a ∼70 basepair sequence that was predicted to form an extended stem loop structure (Figure 5a). This region included a 25 basepair sequence containing a pseudo-palindromic sequence located at the tip of that predicted structure, that overlapped with the -35 element within a predicted bacterial promoter region (Figure 5b) suggested by the BPROM server (49). We therefore tested a ∼200 basepair DNA sequence containing this region in additional EMSAs and observed much higher affinity binding and formation of a shifted protein-DNA complex, indicative of sequence-specific interaction (Figure 5**, inset)**. Titration of the BrxR protein indicated a dissociation constant of approximately 100 nM. Additional experiments, in which flanking basepairs were systematically truncated from either end of the DNA probe, further confirmed the BrxR target site as corresponding roughly to basepairs -51 to -27 upstream of the BREX transcription start site (Supplementary Figure S7).

**Figure 5.**
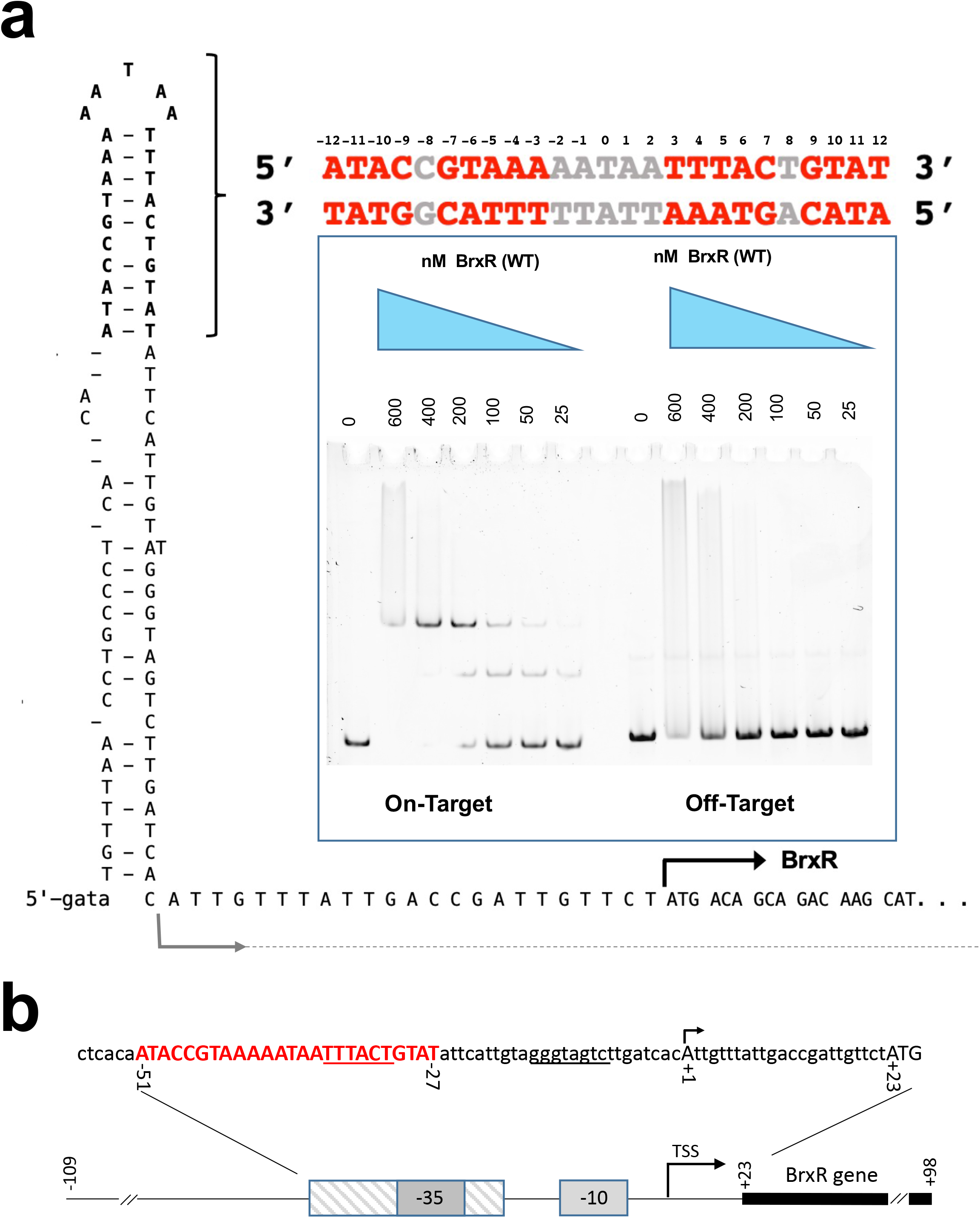
Sequence, predicted secondary structure and BrxR binding site upstream of the BREX operon. See also Supplementary Figure S7. ***Panel a:*** The *Acinetobacter* genomic sequence immediately upstream of BrxR ORF, spanning approximately 200 basepairs, contains an ∼70 basepair sequence that is predicted to form an extended stem loop structure, including a 25 basepair sequence corresponding to a pseudo-palindrome located at the tip of the predicted hairpin (indicated by bracket). When presented to BrxR as a double-stranded DNA construct (red and grey bases), electrophoretic mobility shift binding assays demonstrate binding with a dissociation constant of approximately 100 nM and a subsequent super-shift at higher protein concentrations. The imperfect palindrome formed by the sequence is conserved between half-sites at 9 out of 10 positions, with the half-sites spaced 5 nucleotides apart. The grey arrow illustrates the transcription start site for the operon; the black arrow depicts the translational start site for BrxR. ***Panel b:*** The BrxR binding site (red capitalized bases) overlaps with a predicted bacterial promoter (identified using the BPROM server (49)) that contains canonical -35 and -10 box motifs (underlined bases). The BrxR binding site corresponds approximately to basepairs -51 to -27 (relative to the BREX transcription start site; see also Figure 2 and Supplementary Figure S2). The BREX transcription start site at position +1 is indicated with the arrow; the BrxR translation start codon is indicated by the capitalized ATG.

The gel shift experiments consistently and clearly indicated an initial shift at lower BrxR concentrations to a single slower migrating species, and then a subsequent shift to an even slower migrating species at higher concentrations of protein. This behavior is observed for DNA probes of various lengths and truncations near the defined boundaries of the identified target site, as well as for DNA probes in which the target site is flanked by fully randomized DNA sequence. We believe the simplest explanation for this behavior may be association of DNA-bound protein homodimers into a high order assemblage (such as a dimer of DNA-bound dimers) at elevated protein concentrations.

We next co-crystallized of BrxR with this palindromic sequence and solved a second structure of BrxR bound to the DNA target at 2.3 Å resolution (Figure 6 and Table 1). The cocrystal structure illustrates that BrxR binds the DNA target through extensive interactions between each subunit’s wHTH domain and each corresponding target half-site. The DNA duplex maintains a B-form topology and all the bases are found in Watson-Crick interactions with their partners from the opposing complementary strand. The DNA is bent by approximately 10 to 15° within the plane of the protein-DNA complex, and the minor groove is significantly narrowed across the central five base pair sequence that separates the two related target half-sites. Overlay of the DNA-bound protein structure to the unbound form of BrxR shows a near perfect match in structure (Supplementary Figure S8), indicating that the unbound form of BrxR is in the proper configuration for binding this sequence.

**Figure 6.**
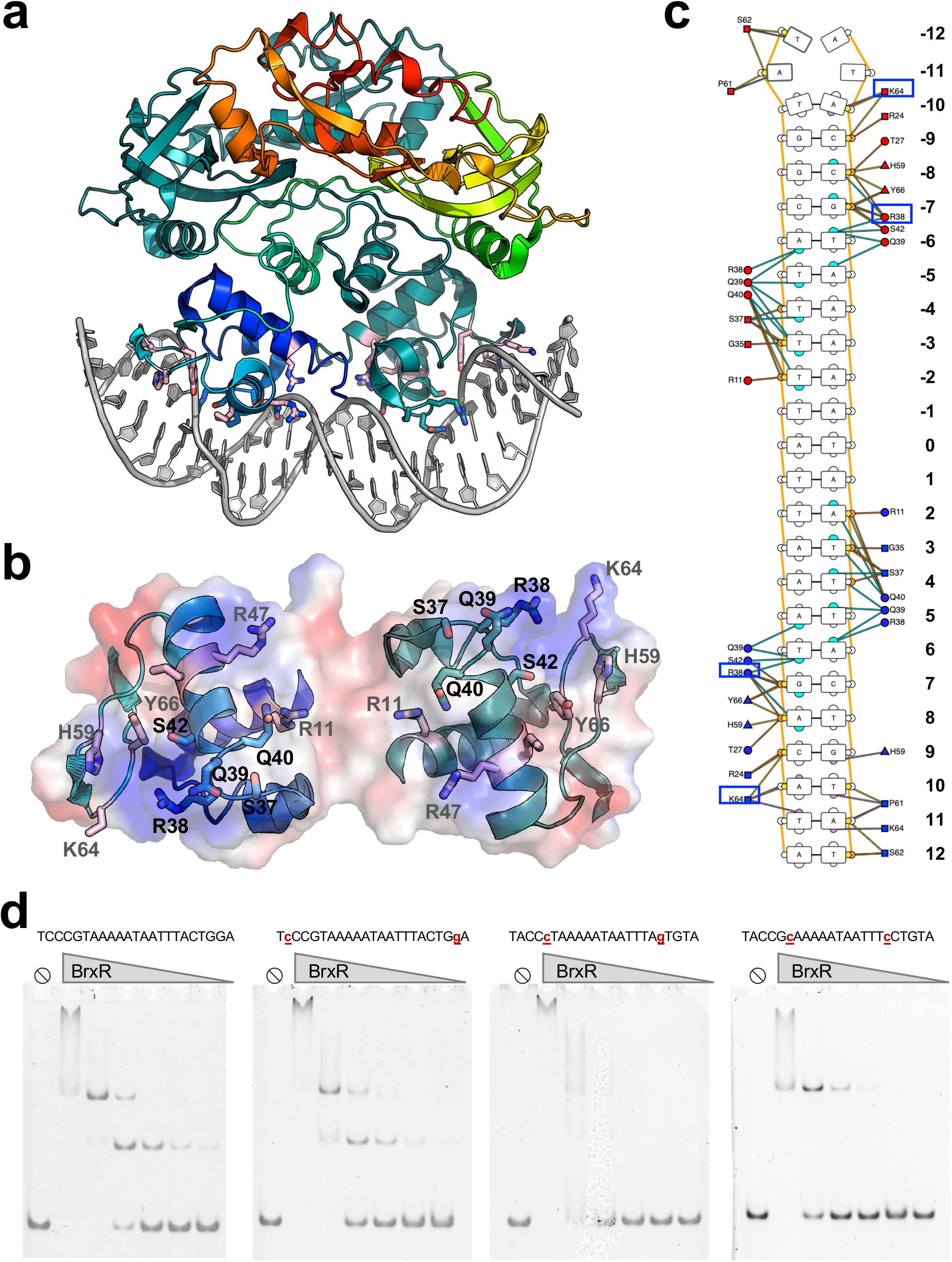
Crystal structure of BrxR bound to DNA target site. ***Panel a:*** Structure of the BrxR homodimer bound to the DNA target shown in Figure 5. The conformation of the DNA-bound protein is closely superimposable with the unbound protein (Supplementary Figure S8) with an overall rmsd across all *α*-carbons of approximately 0.8 Å, indicating that the apo-protein is in the proper conformation to interact with this target sequence. ***Panel b***: The DNA contacting surface formed by each protein subunit’s wHTH domain. ***Panel c:*** Contacts between BrxR and the bound DNA target site extend across 11 bases for each DNA half-site. Base-specific contacts include a group of positively charged and polar residues spanning S37, R38, Q39, Q40, which are part of a conserved SRQQ sequence motif conserved across the closest homologues of *Acinetobacter* BrxR; additional basic residues that contribute include R11, K46, R47, K64, H59 and Y66. R38 (which makes base-specific contacts at positions +/- 6 and +/-7) and K64)which contacts bases in the minor groove at positions +/- 10 and +/- 11) are boxed. Figure panel generated using the DNAProDB online analysis tool (50). ***Panel d:*** Gel shift analyses using BrxR’s 25 bp target sequence flanked by non-specific DNA (total length of the sequences used in these assays is 99 basepairs). The leftmost panel is BrxR’s wildtype target sequence. The substrates in the right three panels contain the indicated substitutions at equivalent positions in both half sites. For each binding assay, BrxR protein was used at 800, 400, 200, 100, 50 and 25 nM. Basepair substitutions at positions +/- 6 or 7 qualitatively reduce BrxR binding affinity (third panel) while basepair substitutions at positions +/-10 display a negligible effect on binding.

Each protein subunit contributes an interface (Figure 6b) that buries approximately 870 Å^2^ of surface area, via the formation direct and water-mediated atomic contacts to the DNA (analysis facilitated and visualized with the DNAProDB online tool (50)). BrxR makes relatively few direct contacts to bases in each half-site of the DNA target. One residue in particular (R38) appears to be involved in direct recognition of two consecutive bases in each half-site (G:C and T:A at positions +/- 6 and 7) (Figure 6c). K64 also appears to make multiple direct contacts, albeit in the minor groove, contacting bases at position +/- 10 and 11. Beyond these direct contacts, the substantial bend in the DNA target likely contributes to specificity across the center of the target, by promoting the A:T rich sequence across those basepairs.

We went on to further validate the observed contacts from the DNA-bound BrxR structure and its specificity by conducting additional gel shift assays using DNA probes in which the target site harbored mutations at the basepair positions involved in direct contacts with protein side chains described above, or in which the spacer between the target half-sites was altered by a single basepair (i.e. from N_5_ to N_4_) (Figure 6d). In these experiments, the target sites were flanked by identical, fully randomized DNA sequences. Basepair substitutions at positions +/- 6 or 7 (contacted by R38) qualitatively reduced BrxR binding affinity, while basepair substitutions at positions +/-10 (contacted by K64) or reduction in the spacer length by one basepair had a negligible effect on binding.

R38 is part of series of four sequential residues (S_37_-R_38_-Q_39_-Q_40_) that are highly conserved among many BrxR_Acin_ homologues Similarly, K64 is part of series of three sequential residues (K_64_-G_65_-Y_66_) that are also conserved across the same homologues (Figure 7). This suggests that that those BrxR proteins may bind to similar DNA targets in other organisms. In contrast, the same residues in the more distantly related *E. fergusonii* BrxR protein (final sequence in Figure 7), which has also been recently characterized (51) are not conserved (’APSVAT’ rather than ‘SRQQ’ and ‘RVH’ rather than ‘KGY’), presumably corresponding to the recognition of a more distantly related target sequence (51).

**Figure 7.**
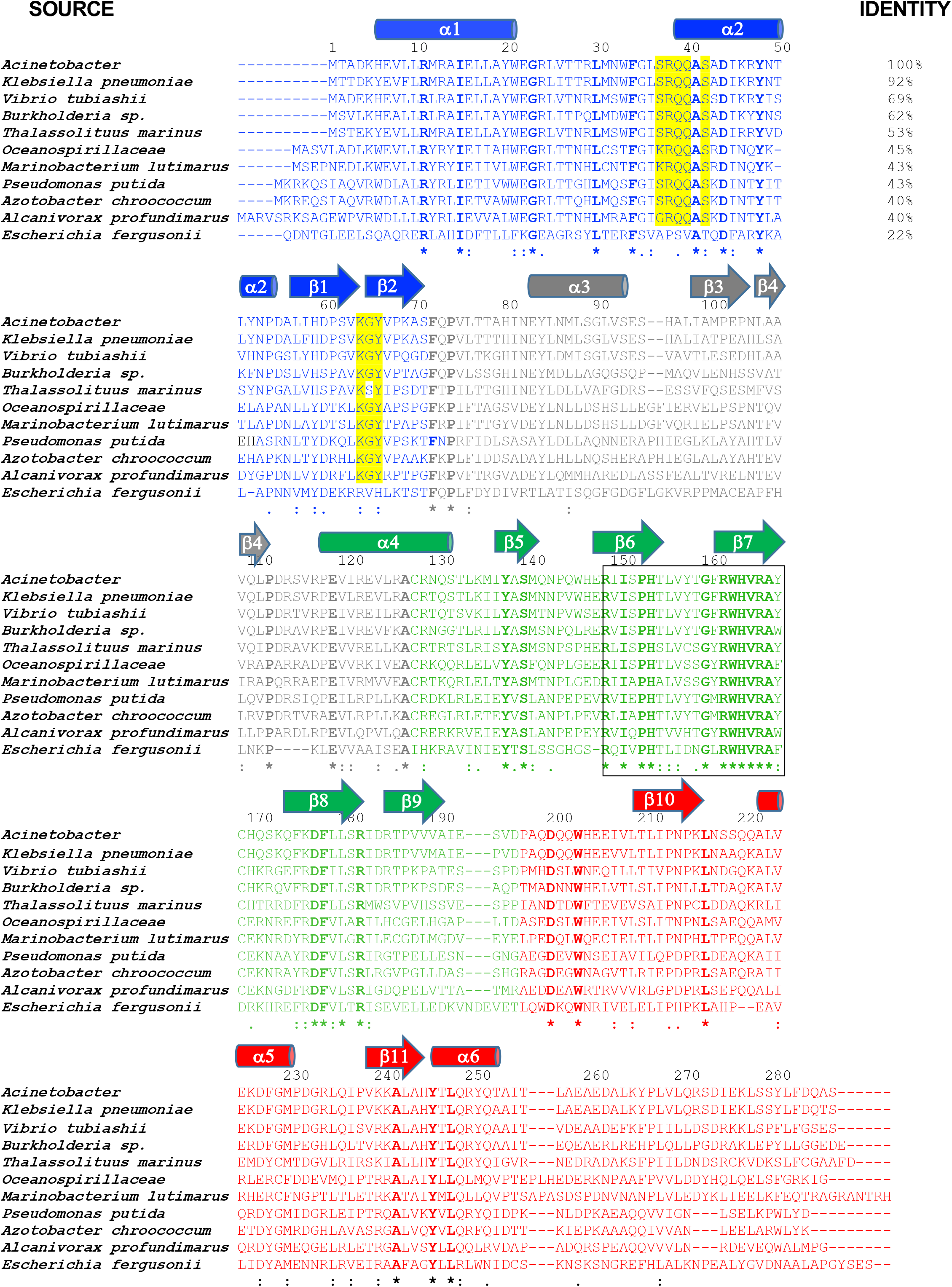
Multi-sequence alignment of BrxR homologues. Residue coloring indicates the domains of the protein as similarly colored in Figure 4. Residues that are entirely conserved across a wide range of identities relative to BrxR_Acin_ (22% overall identity) are indicated with bold font and asterisks. Residues that make contacts to individual DNA bases in the target site (Figure 6) and that are conserved for homologues down to approximately 40% overall sequence identity are highlighted yellow. Residues that comprise the conserved core of the WYL domain are indicated with a box (also indicated in Figure 4).

### Deletion of the BrxR gene affects bacterial growth and phage restriction differently than elimination of the BrxR protein itself

Based on the structural and biochemical analyses described above, we generated two additional disruptions to BrxR in the pACYC184-BREX vector and validated them by sequencing across the entire plasmid: a premature stop codon (created by altering TGGg into TAGg via a ‘G’ to ‘A’ substitution in codon 21 of the BrxR gene; the nucleotide after the stop codon is indicated in lower case) or a point mutation (R47A) in the helix-turn-helix DNA binding surface that was predicted to interfere with DNA binding. The R47A amino acid substitution was also incorporated into the recombinant protein expression vector, and the corresponding construct was purified (Figure 8a) and examined to ensure that the mutation still resulted in soluble homodimeric protein (Figure 8b), did not cause a significant destabilization of the protein fold (Figure 8c), and eliminated site-specific DNA binding as predicted (Figure 8d). As a further control for experiments examining the effect of R47A on BrxR behavior and function, several additional point mutations were introduced into the WYL domain of the protein; the resulting constructs were individually purified, and their solution behavior and stability characterized (Supplementary Figure S9a-d). One construct from that series (R149A, which is found within a conserved basic pocket in the protein’s WYL domain; see Figure 4e) was shown to still bind the protein’s target site similarly to the wild-type protein (Supplementary Figure S9e).

**Figure 8.**
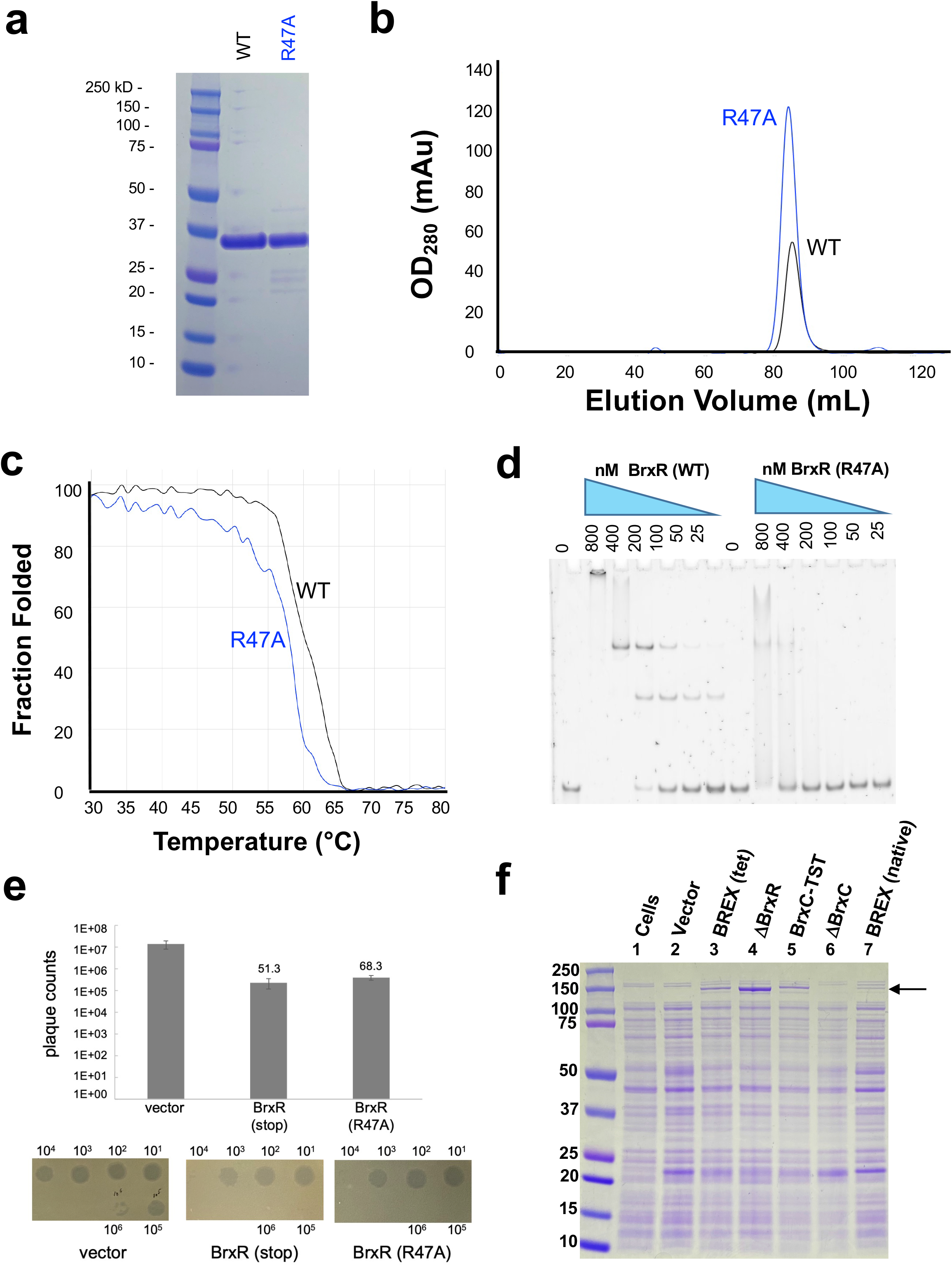
Characterization of a BrxR point mutant with reduced DNA target binding affinity. ***Panels a-c***: BrxR containing a single point mutation in its DNA binding domain (R47A) was expressed and purified to homogeneity; it behaves similarly to the wild-type protein dimer in solution as indicated by its elution profile on size exclusion chromatography (SEC) and displays similar thermal stability and unfolding behavior as the wild-type protein. See also Supplementary Figure S9. ***Panel d***: Electrophoretic mobility shift assays (EMSA) demonstrate a significant reduction in DNA target binding by purified BrxR (R47A) protein. ***Panel e:*** Phage restriction plaque assays indicate that unlike the effect of a precise deletion of the BrxR gene (Figure 3; which causes significant toxicity and reduction in BREX restriction activity), the disruption of the BrxR gene by incorporation of an early stop codon or by introduction of a mutation that blocks BrxR DNA binding activity has little effect on cell growth or phage restriction in the same assays. ***Panel f:*** SDS PAGE analysis of whole cell lysates of *E. coli* ER2683 cells transformed with the following pACYC-BREX constructs: cells only (lane 1); pACYC vector only (lane 2); WT BREX (lane 3); a precise deletion of the BrxR gene (ΔBrxR; lane 4); an epitope-tagged version of BrxC that runs slightly higher than untagged BrxC (BrxC-TST; lane 5); a precise deletion of the BrxC gene (ΔBrxC; lane 6); BREX expressed from its native promoter (lane 7). Transformation with pACYC harboring wild-type BREX results in the appearance of a novel band at approximately 140 to 150 kD (arrow), and deletion of the BrxR gene results in significant increase in its intensity (compare lanes 3 and 4). The same band disappears altogether when BrxC gene is deleted from the BREX operon (lane 6) and shifts slightly upward when BrxC is fused to a 35 amino acid twin-strep tag (lane 5), indicating that the upregulated protein product is BrxC. This same band is present in when the BREX operon is expressed from the native promoter (lane 7), albeit at a reduced level compared to when BREX is expressed from the constitutive tet promoter (lane 3).

Whereas the precise deletion of the BrxR gene from the BREX operon significantly reduced phage restriction (Figure 3 and Supplementary Figure S4), cells transformed with the BREX operon harboring either the premature stop at codon 21 or an R47A point mutation in the BrxR gene displayed similar abilities to restrict *λ*_vir_ in plaque formation assays (Figure 8e).

Finally, we examined protein expression in *E. coli* cells in the presence and absence of the BREX operon, which itself harbored either an intact copy of the BrxR coding sequence or a deletion or a disruption of that gene. Polyacrylamide gel electrophoresis (PAGE) analyses (Figure 8f) indicated the visible expression of at least one additional protein from cells harboring the wild-type BREX operon, corresponding to a molecular weight similar to BrxC or PglX. The expression of that protein was significantly increased in cells that contained a precise deletion of the BrxR gene. Additional control lanes, corresponding to cells that harbored either a clean deletion of BrxC or an epitope-tagged version of BrxC, validated that protein as the BREX factor being significantly upregulated in the absence of the BrxR gene. In contrast, cells harboring the BREX operon with a premature stop codon in BrxR displayed little difference in protein expression levels as compared to those harboring the wild-type BREX operon. This appears to reflect our earlier observation that disruption of the BrxR gene and eliminating BrxR protein expression via the same stop codon (rather than precisely deleting the gene) had little measurable effect on phage restriction.

## DISCUSSION

### The distribution and diversity of WYL proteins within bacterial defense islands and BREX systems is widespread

A previously described bioinformatic analysis of PafBC, a bacterial transcriptional regulator involved in DNA damage response that contains a similar HTH-WYL-WCX domain organization as BrxR, identified over 10,000 homologues that are distributed across actinobacteria, proteobacteria, firmicutes and bacterioidetes (25). Similarly, our own BLASTP search (52) using the *Acinetobacter* 394 BrxR sequence (‘BrxR_Acin_’) as a starting query returns over 9000 hits with sequence identities of approximately 20% or higher (corresponding to E-values of 28 or better). BrxR homologs spanning a wide range of sequence identities (as low as 25%) are frequently linked to defense systems across a wide range of bacterial phyla, indicating a strong selective pressure to maintain BrxR association with these islands. Many of those BrxR homologues are found within gamma proteobacteria (which includes enterobacteria such as *Escherichia fergusonii*, Vibrionaceae including *Vibrio cholera*, and Pseudomonacaea including the *Pseudomonas* genera). However, there are numerous BrxR homologues found in alternative bacterial clades and genera, which one might expect for a regulatory protein frequently associated with mobile genetic islands and their corresponding phage defense and antibiotic resistance systems. A recent parallel study of BrxR conducted using a BREX system from *Escherichia fergusonii* (51) has indicated that nearly half of identifiable BrxR proteins are associated with a wide variety of known phage defense systems, both in isolation and in the context of genetic defense islands. Yet another parallel study (53) has further demonstrated that a similar WYL regulatory protein, termed ‘CapW’, is involved in control of a CBASS phage defense system, with structural and mechanistic features that are similar to those reported in this study and in (51). Collectively, these three studies indicate that BrxR/CapW represent a broad superfamily of proteins involved in regulation of phage defense.

We constructed a multi-sequence alignment (Figure 7) from a sampling of nine BrxR sequences spanning amino acid identities, relative to BrxR_Acin_, ranging from 92% (*Klebsiella*) to 22% (*Escherichia fergesonii*, which has also been recently characterized (51)). Residues involved in contacting individual base pairs in the BrxR_Acin_ DNA target site (in particular, those spanning a motif corresponding to S_37_R_38_Q_39_Q_40_ in BrxR_Acin_; Figure 4) are conserved among homologues that exhibit down to 40% amino acid identity to BrxR_Acin_. In comparison, BrxR from *Escherichia fergusonii* (which recognizes a different pseudo-palindromic target site in the same region upstream of its own BREX operon) is diverged across those same residues. Overall, 64 residues are strictly conserved across the entire range of homologues shown in our alignment; of those, a cluster of residues correspond to a conserved basic pocket in the WYL domain (illustrated in Figure 6).

### The BrxR gene and BrxR protein appear to play unique cis- and trans-regulatory roles

The observation that the effect of a precise deletion of the BrxR coding sequence from the BREX operon (which resulted in toxicity and a reduction in phage restriction) differ from the negligible effect of an early stop codon in the BrxR coding sequence (that otherwise left the BrxR gene intact) was unexpected. This result was reproduced in a separate experiment in which an amino acid mutation (R47A) that abrogated BrxR binding to its DNA target, was instead introduced into the BREX operon. These results imply that in the Acinetobacter 394 BREX system, the BrxR gene itself may contain additional *cis* regulatory elements, such as an alternative promoter, that when removed disrupt the normal expression of BREX ORF’s, and in turn the normal restriction function of BREX (Figure 3). Consistent with this possibility, an analysis of the BrxR coding sequence from *Acinetobacter* using the BPROM server (49) suggests the possible presence of a promoter within the BrxR gene.

The toxicity of E. coli harboring the BREX operon with a precise deletion of the BrxR ORF (‘ΔBrxR’), as well as those with deletions of the BrxB or PglZ genes, may be caused by an imbalance of proteins that form toxin-antitoxin pairs. PglX and PglZ have previously been implicated as toxin-antitoxin partners, and it has been postulated that BREX may contain additional similar interactions (17). The marked increase in BrxC protein levels that we observe in ΔBrxR cells relative to cells expressing an intact BREX operon (Figure 8F) may implicate BrxC as being such a toxin (although, if true, we do not know its antitoxin partner).

### Potential regulation of BrxR activity

The observation that BrxR binds to a specific DNA target upstream of the BREX operon at a position that overlaps with a predicted bacterial promoter (Figure 5) indicates that it is likely a transcriptional regulator. However, the presence of a highly conserved region in BrxR’s WYL domain further implies that it may be involved in additional layers of regulation that depend upon the binding of an effector molecule (perhaps a bacterial second messenger or a modified nucleotide derived from a phage genome) that is produced during a phage challenge. The binding of such a molecule might alter the relative binding affinity of BrxR towards competing DNA targets, as has been previously hypothesized for the PafBC regulator as part of a bacterial DNA damage response (24, 25). Unlike PafBC, which appears to require a significant domain rearrangement to bind DNA (thought to be driven by binding of an unknown effector to the WYL domain), the BrxR protein visualized in this study is preorganized in a homodimeric conformation that is appropriate for target binding with negligible conformational changes to the protein architecture. Therefore, any regulated changes in DNA binding specificity for BrxR might be expected to involve relatively subtle structural rearrangements.

The published literature provides a number of anecdotal lines of evidence that proteins containing WYL domains might act as transcriptional regulators that display differential activities (involving alteration of binding activities and specificities, and/or transitioning between repressive or stimulatory functions) as part of a corresponding defense system’s response to a foreign invader or challenge (22). For example, a WYL domain protein (sll7009) appears to act as a transcriptional repressor of a CRISPR-Cas system in *Synechocystis sp* (*23*), and WYL proteins have been shown to regulate other CRISPR systems (54). Two recent studies both demonstrate that WYL domain proteins found in BREX systems associated with highly mobile SXT Integrative and Conjugative Element (‘SXT ICE’) defense islands, in *Vibrio cholerae* and *Proteus mirabilis* respectively, appear to play important regulatory roles in BREX action (20, 21). Our own alignment of WYL domains from these elements (which share ∼ 25% sequence identity with BrxR_Acin_) and subsequent structural modeling carried out with the AlphaFold server (55) (Supplementary Figure S10) indicates that those proteins also share the same wHTH-WYL-WCX domain organization and appear to adopt a conformation of those domains that is seen in our crystal structures of the BrxR_Acin_ homodimer.

The presence of the WYL domain in BrxR_Acin_ might confer a sensitivity to an effector molecule as part of an early response to a phage challenge. If true, it is interesting to speculate on the nature of such an effector and the possibility that such an effector might modulate BrxR regulation of BREX activity. The highly conserved pockets in the WYL domain found across BrxR homologues (Figure 4cde and Figure 8) appear to be positively charged, which could be compatible with binding to a nucleotide-derived ligand. Those pockets are loosely associated with the final few residues of the WCX domain’s C-terminal tail, which might be displaced upon effector binding. As well, two adjoining a-helices in the WCX domains, that are immediately upstream of their C-termini (residues 253-263) partially occlude a considerably longer highly basic channel that connects the conserved pockets in the two WYL domains; if also displaced a surface of sufficient area and dimensions to bind a larger moiety (such as a DNA or RNA oligonucleotide) would be exposed. Consistent with this latter possibility, other proteins have been observed to use WYL and WCX domains to bind DNA or RNA effectors (25,56–60).

We performed experiments to examine the effect of a collection of known bacterial second messengers (including a range of cyclic mono- and dinucleotides) on the affinity of BrxR towards the target described in this study but did not observe a significant effect.

### Potential functions of BrxR as a BREX regulator

The observation that an early stop codon in BrxR does not affect the ability of BREX to restrict *λ*_vir_, coupled with the overlap of the BrxR binding site with a promoter in its upstream region, suggests that BrxR may function as a repressor of the BREX system. Such a role has been proposed for other WYL protein-mediated regulation of defense systems, including BrxR and CapW (20,23,51,53). We speculate there may be either (or both) of two potential roles for an inhibitory function of BrxR:

1. Function as a “2nd line of defense”. Activation of the BREX system in its native context may require release from a repressed state, perhaps mediated by BrxR. However, such a mechanism poses a possible conundrum for the bacteria: if cells were to rely on BREX as a ‘front line’ defense, the time required for the system to emerge from a repressed state might be too slow relative to the initial appearance and action of phage. Perhaps instead, BREX might provide a ‘second line’ of defense that is activated during the time that first responders (such as restriction-modification and CRISPR systems) successfully ward off an initial challenge. The BREX system would then be primed and ready if resistant phage break through these front lines of defense. One might imagine that a rare phage variant with appropriate resistance factors (perhaps present at very low levels in the initial phage population, and then selected for during the first early stages of a challenge) might eventually create a ‘second wave’ requiring additional strategies for restriction.
2. Facilitate horizontal transfer into new hosts. It is notable that in various bacterial systems (such as Vibrio cholera (20), Proteus mirabilis (21) or Escherichia fergusonii (12)), BrxR homologues are coupled to a wide range defense systems (including BREX) that are contained within mobile elements, suggesting that these defense systems frequently transfer between hosts. We postulate that BrxR may transiently suppress a BREX-mediated toxic activity during horizontal gene transfer so that PglX (and associated factors) has time to methylate and protect the new host’s genome. Such a role has been described for C-proteins during horizontal transfer of R-M systems (13, 14).

## ACKNOWLEDGEMENTS

We thank Dr. Harmit Malik for helpful discussions and for support of KF and DH. Work performed by KF and DH was further supported by a grant from the Mathers Foundation and by the HHMI. Support was also provided New England Biolabs and by the NIH for both BLS (R01 GM105691) and BKK (R15 GM140375). The Berkeley Center for Structural Biology is supported in part by the National Institutes of Health, National Institute of General Medical Sciences, and the Howard Hughes Medical Institute. The Advanced Light Source is supported by the Director, Office of Science, Office of Basic Energy Sciences, of the U.S. Department of Energy under Contract No. DE-AC02-05CH11231. The Pilatus detector was funded under NIH grant S10OD021832. The ALS-ENABLE beamlines are supported in part by the National Institutes of Health, National Institute of General Medical Sciences, grant P30 GM124169.

## DATA AVAILABILITY

The sequence of the entire BREX operon is provided as supplementary material for easy download and also is deposited at at Genbank with protein ID WP_176538600.1. The *Acinetobacter* assembly accession ID is ASM1337479v1, leading to all 15 genome sequences (chromosome (CP055277.1) and 14 plasmids (CP055278.1 through CP055291.1). All sequencing data is deposited in the Sequence Read Archive (SRA). The *Acinetobacter* genome sequencing BioProject number is PRJNA638470 (Biosample number SAMN15195663; txid number txid2743575). The *E. coli* methylomics sequencing BioProject number is PRJNA8124724. The transcriptomic TSS sequencing BioProject number is PRJNA814726. The original source data and raw images corresponding to the biochemical analyses of BREX and BrxR function and activity have been uploaded to the Harvard Dataverse public repository (https://dataverse.harvard.edu/dataverse/BrxR). The crystallographic structures described in this manuscript have been deposited in the RCSB protein database (PDB ID codes 7T8K and 7T8L).

## SUPPLEMENTARY FIGURES

**Supplementary Figure S1.**
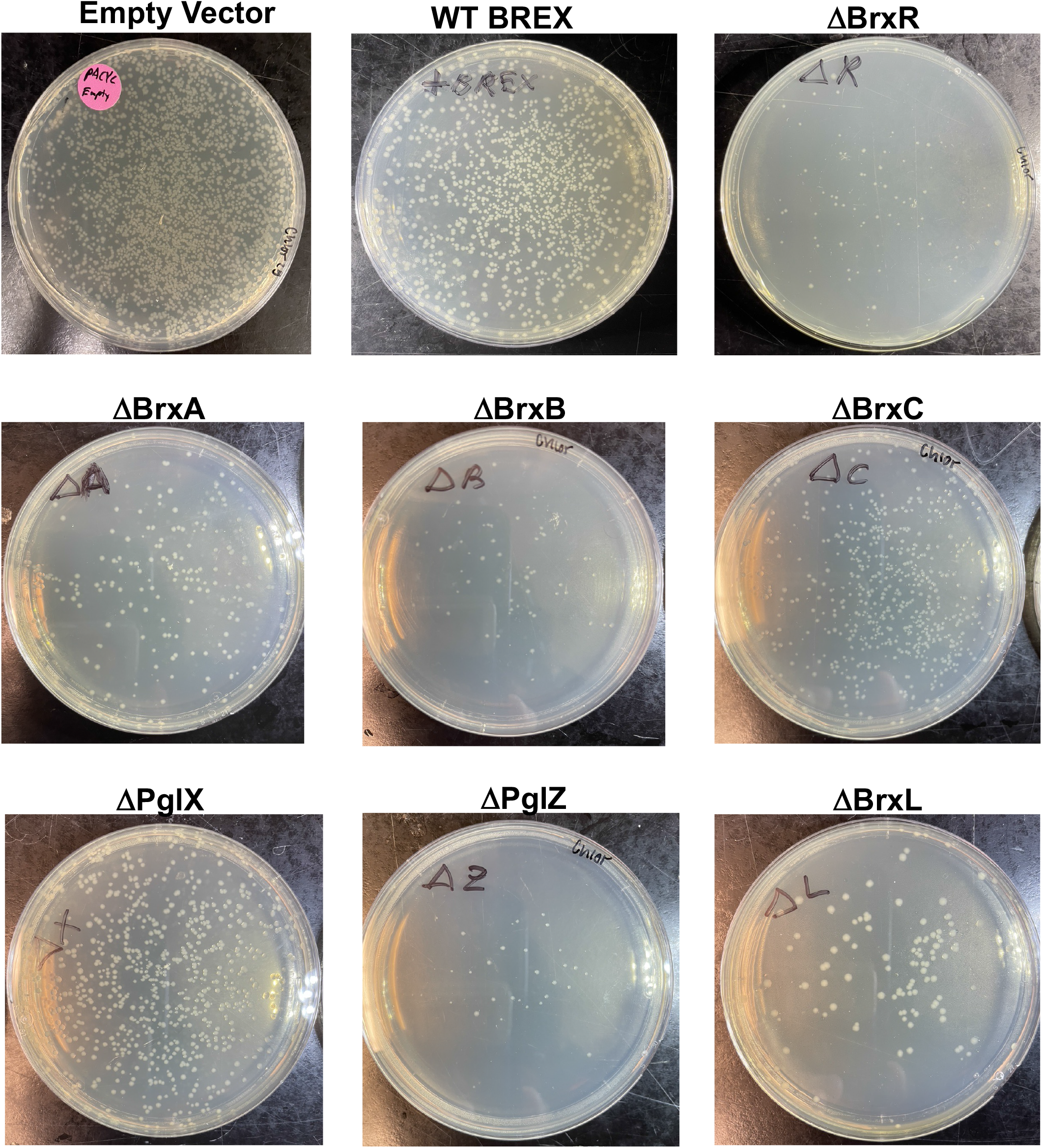
*E. coli* ER2683 cells transformed with empty vector, vector encoding the WT BREX operon, or the same operon harboring precise deletions of each BREX protein factor. Reduced colony size and transformation efficiency is observed for ΔBrxL, ΔBrxB and ΔPglZ.

**Supplementary Figure S2.**
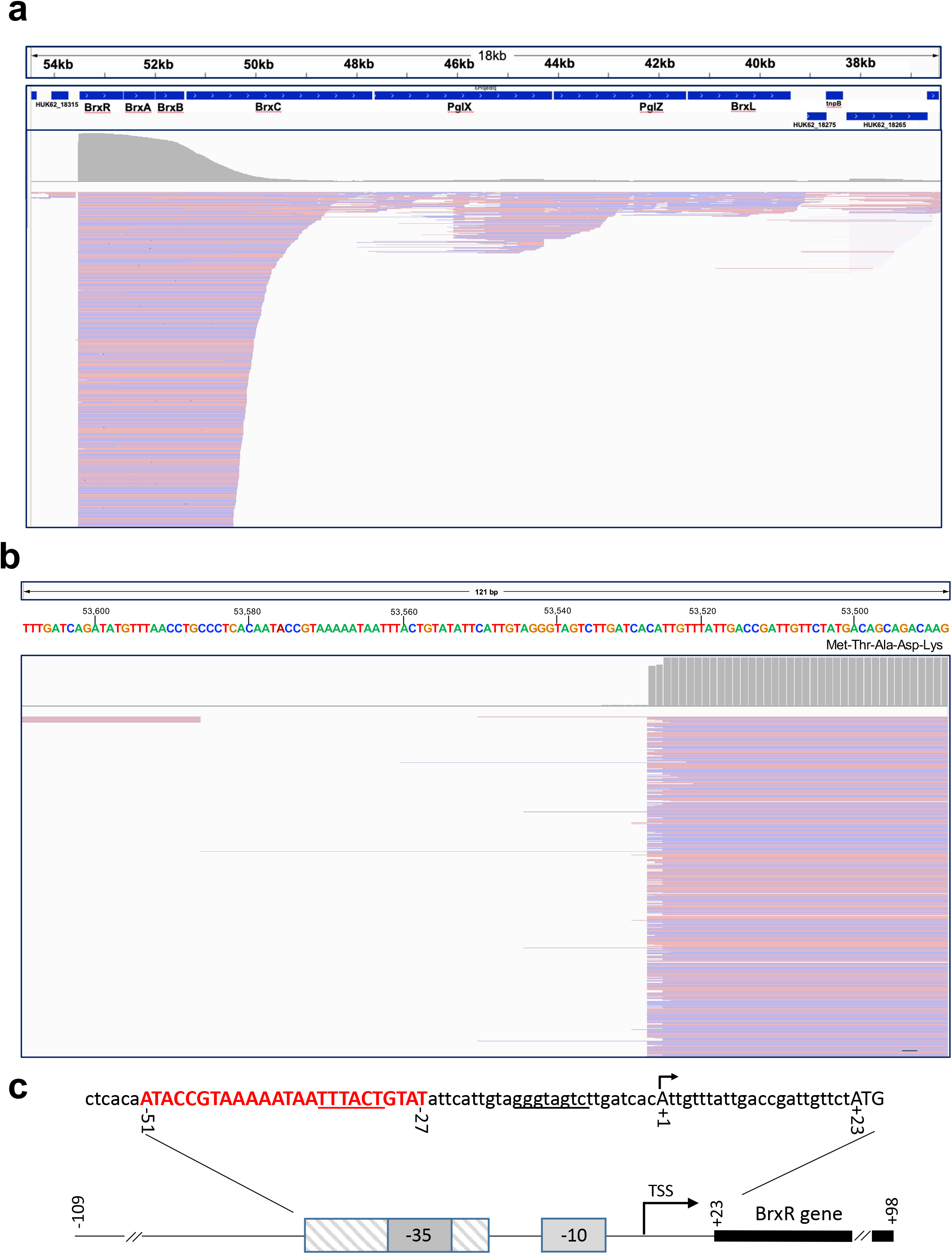
Transcriptional start site (TSS) analysis of *Acinetobacter sp* NEB394 BREX operon. ***Panel a:*** Integrative Genomics Viewer (IGV) representation mapping PacBio SMRT-cappable-seq transcriptional start site data to the Acinetobacter BREX region in plasmid pBspH3. The *top track* shows position within the pBspH3 plasmid and the *second track* shows the BREX reading frames. The *third track* shows read coverage (gray, scale 0 to 3000) and *bottom track* shows mapped reads (scale 1 to 1000; note PacBio read strand output is arbitrary: reads in salmon map to forward strand, reads in blue map to reverse strand). This shows a strong TSS just before BrxR and the BREX operon, with only very low amounts of internal operon reads at various start positions, indicating one TSS for the entire operon. ***Panel b.*** SMRT-cappable-seq with a detailed view of the region upstream of the BrxR reading frame and the corresponding transcription start site. ***Panel c.*** Sequence of the Acinetobacter sp NEB394 BREX 5’ UTR, indicating the location of the BrxR binding site (red, capitalized), predicted promoter region (underlined), transcription start site (+1, black arrow) and the start codon for the BrxR gene (‘ATG’ at +23).The same figure panel also used in Figure 5b.

**Supplementary Figure S3.**
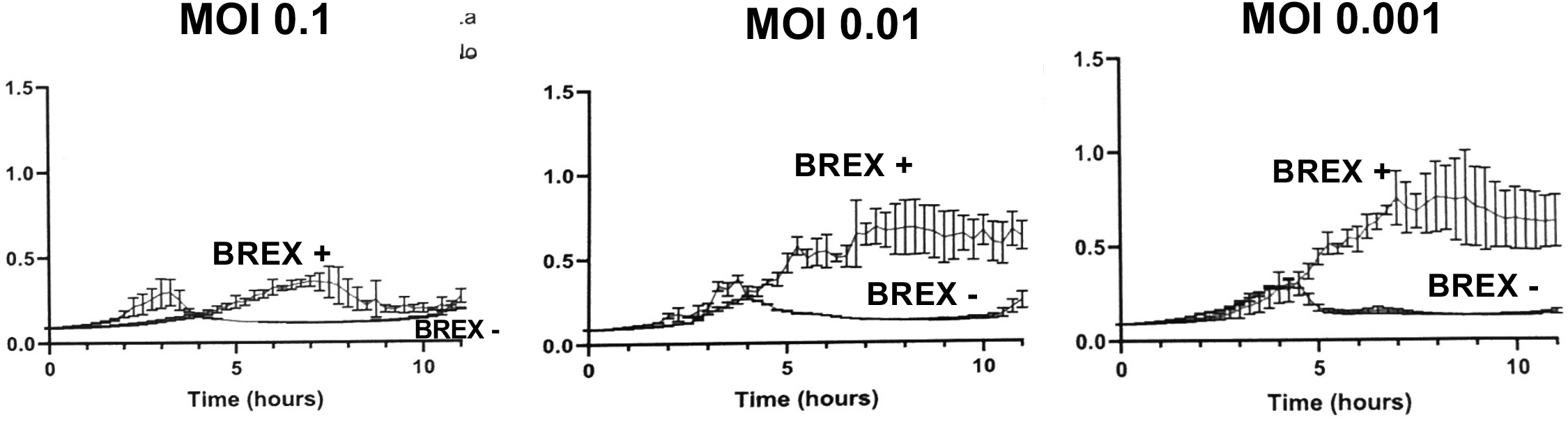
Phage restriction in *E. coli* NEB-5*α* by BREX. ***Panel a:*** Growth of *E. coli strain* NEB5*α* (New England Biolabs) transformed with pACYC184-BREX and challenged with λ_vir_ phage at MOI’s ranging from 0.1 to 0.001. BREX^-^ cells display lysis within five hours of the challenge, versus continued growth and saturation of the cells that harbor the BREX system. The level of protection conferred by the intact BREX system displayed a dependence on the phage MOI; cells containing BREX eventually crashed at the highest MOI (0.1) but continued to grow at MOI’s of 0.01 and 0.001.

**Supplementary Figure S4.**
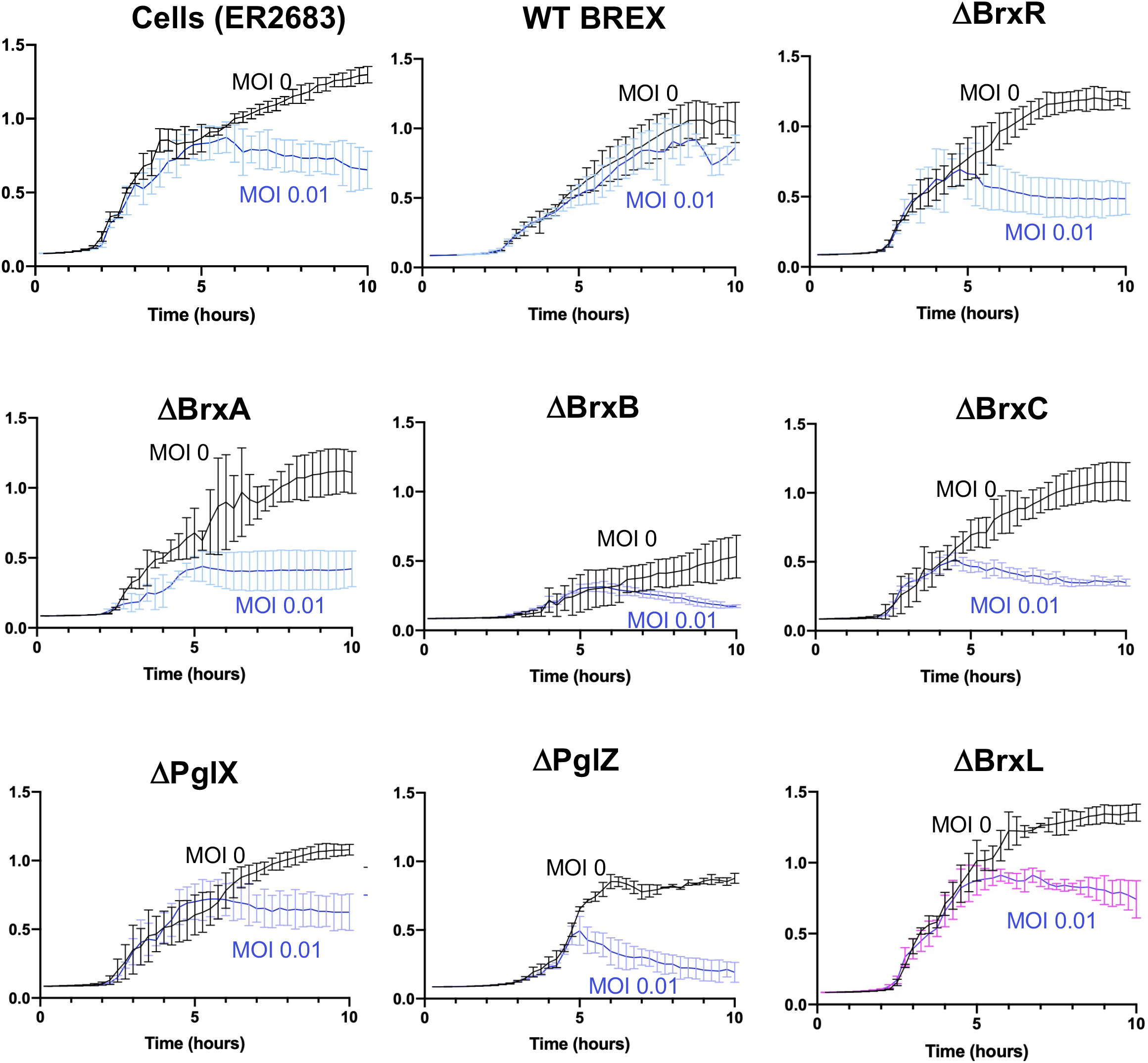
*E. coli* strain ER2683 transformed with pACYC-BREX constructs containing the indicated single deletions were challenged with λ_vir_ phage at an MOI of 0.01. All BREX ORF’s were required for restriction.

**Supplementary Figure S5.**
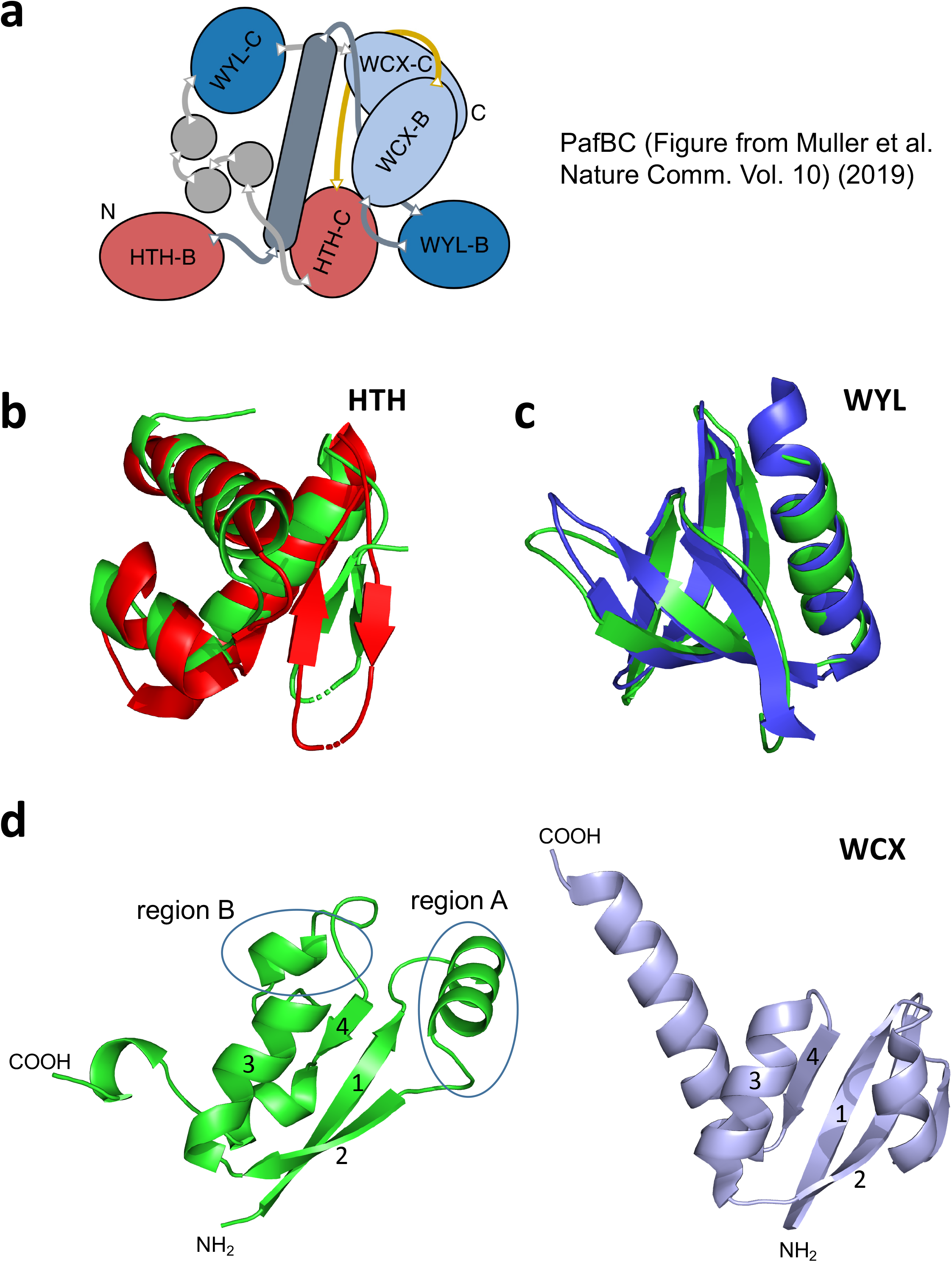
Comparison of BrxR and PafBC structures. ***Panel a:*** PafBC comprises a tandem duplication of wHTH, WYL and WCX domains on the same polypeptide. The unbound form of PafBC (shown here) lacks the 2-fold symmetry present in the homodimeric structure of BrxR. Figure from Muller, 2019 (2). ***Panel b***: Overlay of PafBC’s HTH-B domain with BrxR residues 1-71. ***Panel c:*** Overlay of PafBC’s WYL-C domain with BrxR residues 119-193. ***Panel d:*** BrxR residues 211-291 aligned with PafBC’s WYL-C domain; the domains are shown side-by-side for clarity. The domains share the same core topology (beta-beta-alpha-beta), indicated by numbering of secondary elements on the respective domains. Regions A and B indicate elaborations on this core fold. In Region A (formed between beta(1) and beta(2)), BrxR contains an a-helix, whereas PafBC contains a b-strand and a-helix. In Region B (formed between helix(3) and beta(4), BrxR contains an a-helix, whereas PafBC contains a short linker.

**Supplementary Figure S6.**
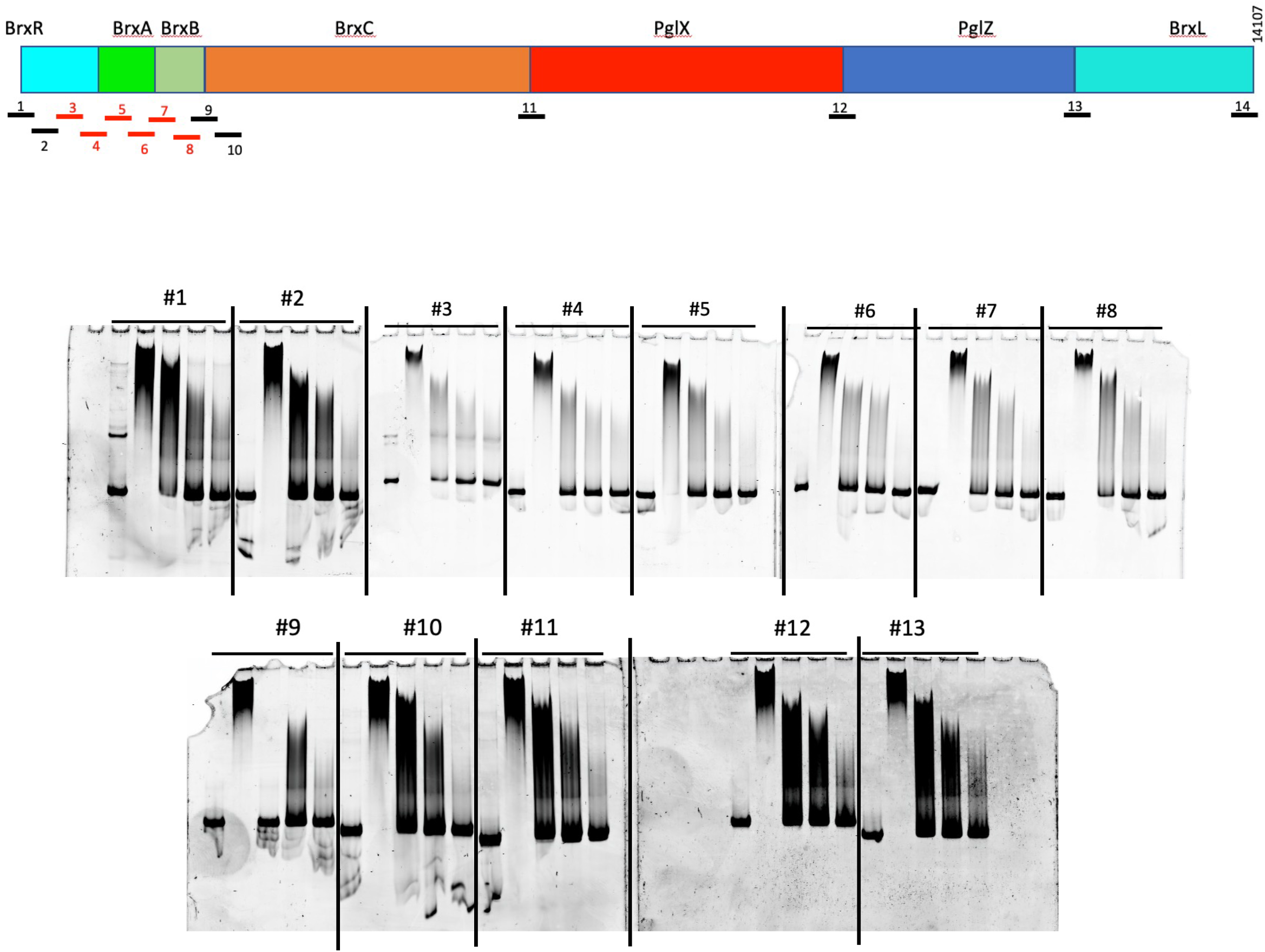
Electrophoretic mobility (‘gel shift’) analyses of BrxR interaction with 13 regions distributed across the BREX operon (positions indicated in top schematic). For each experiment (which are distributed across multiple gels in a composite figure below) lanes with a constant 10 nM concentration of dsDNA probe are incubated with 0 nM BrxR (left-most lane) and then a decreasing titration series of BrxR concentrations corresponding to 800, 400, 200 and 100 nM BrxR. In all cases, no observable specific binding is observed at 100 nM BrxR, and increased concentrations produce a non-specific smear of shifted DNA species.

**Supplementary Figure S7.**
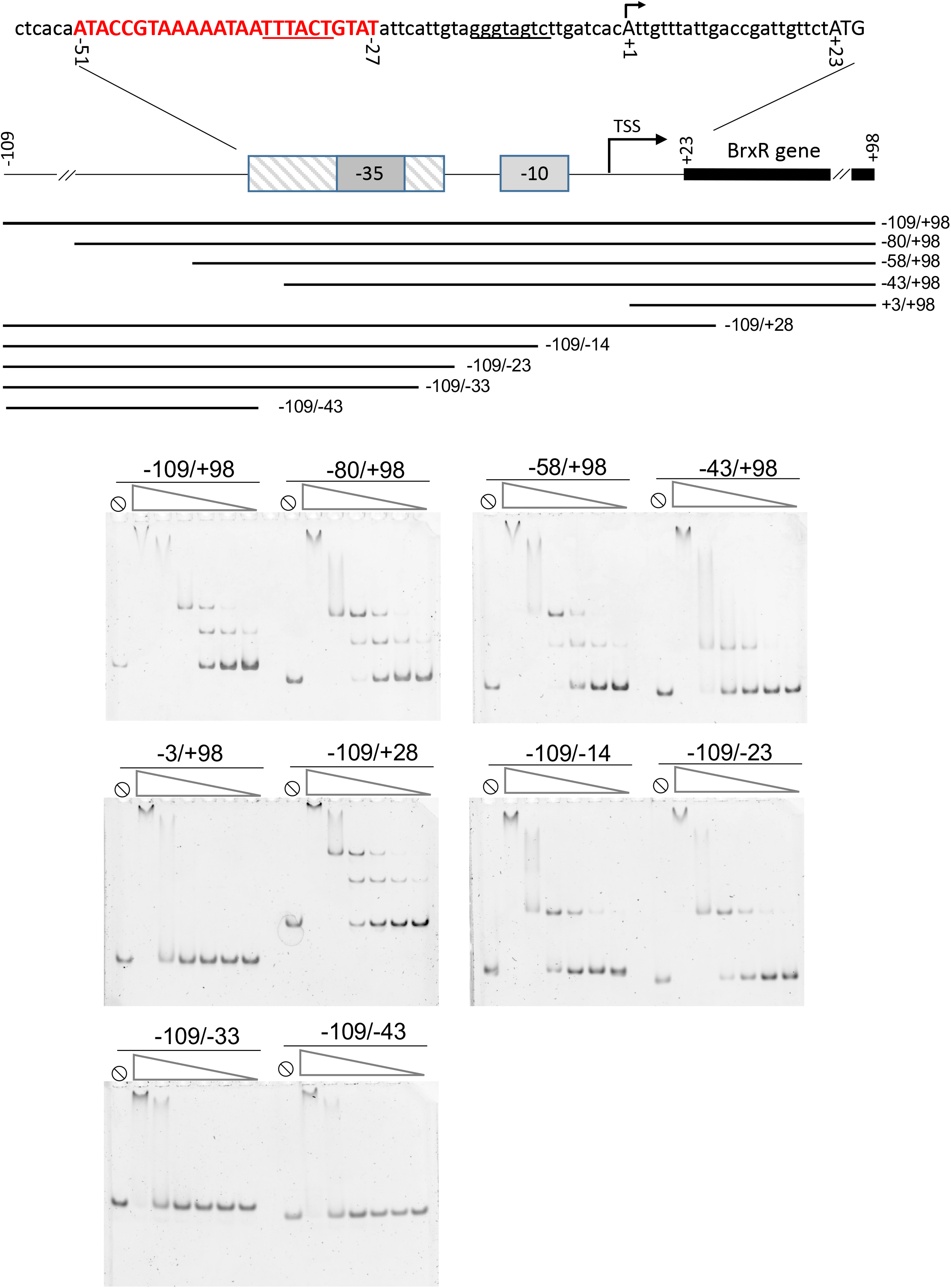
Electrophoretic mobility (‘gel shift’) analyses of BrxR interaction with various DNA probes spanning portions of the 5’ untranslated region preceding the BREX operon. The relative truncations sites for the DNA probes used in the analysis are indicated below the schematic of the 5’ UTR region. The analysis indicates the BrxR binding site (red uppercase font) corresponds approximately to basepairs -51 to -27 (relative to the BREX transcription start site; see Supplementary Figure S2). The -35 and -10 elements of a predicted bacterial promoter (the former of which overlaps with the BrxR binding site) are indicated by <u>underlined bases</u>. The BREX transcription start site at position +1 is indicated with the arrow; the BrxR translation start codon is indicated by the capitalized ATG.

**Supplementary Figure S8.**
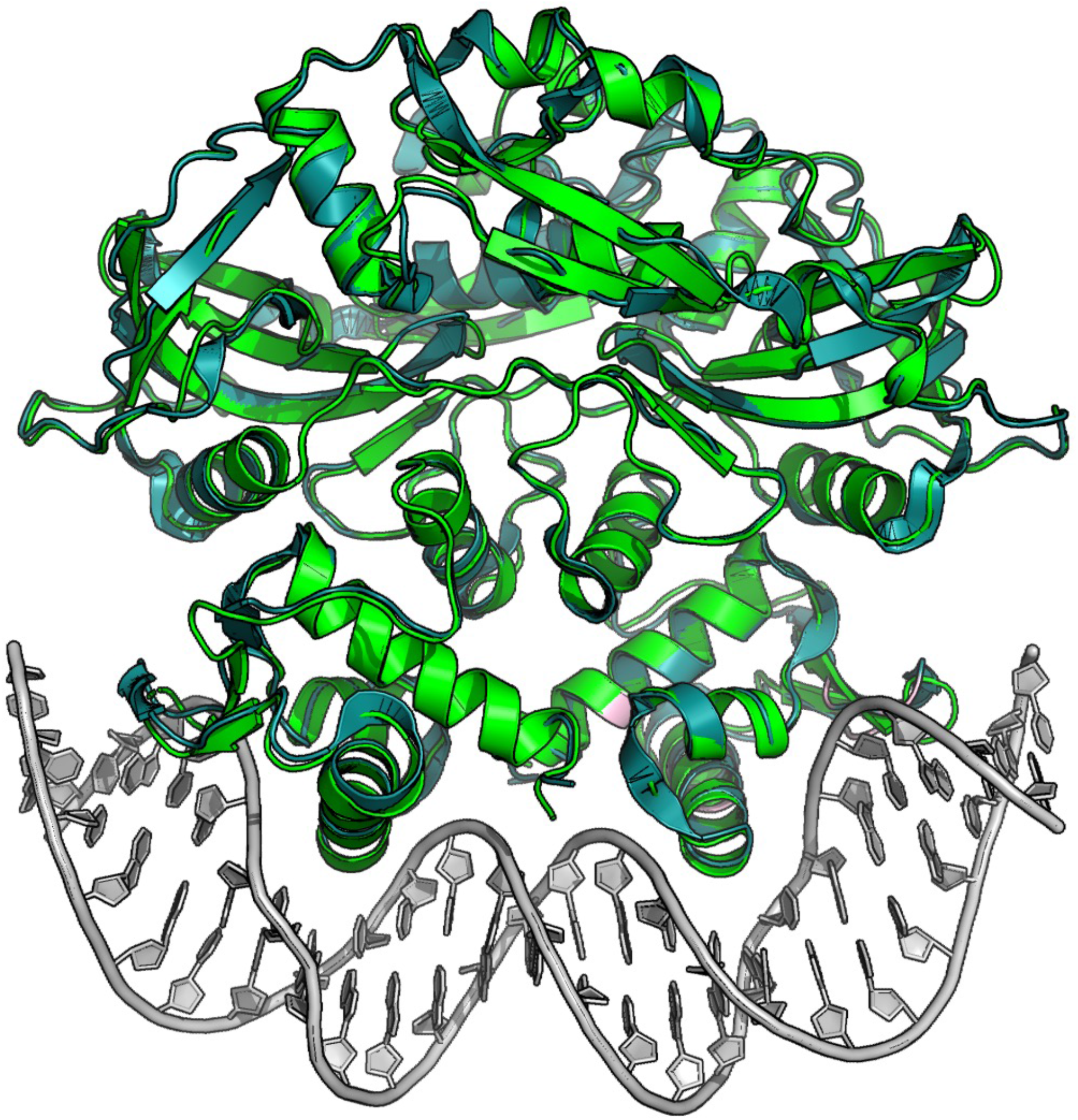
Overlay of the apo-(cyan) and DNA-bound forms of BrxR shows that the protein undergoes very minor conformational changes upon binding DNA. The two forms share an RMSD across all *α*-carbons of approximately 0.8 Å.

**Supplementary Figure S9.**
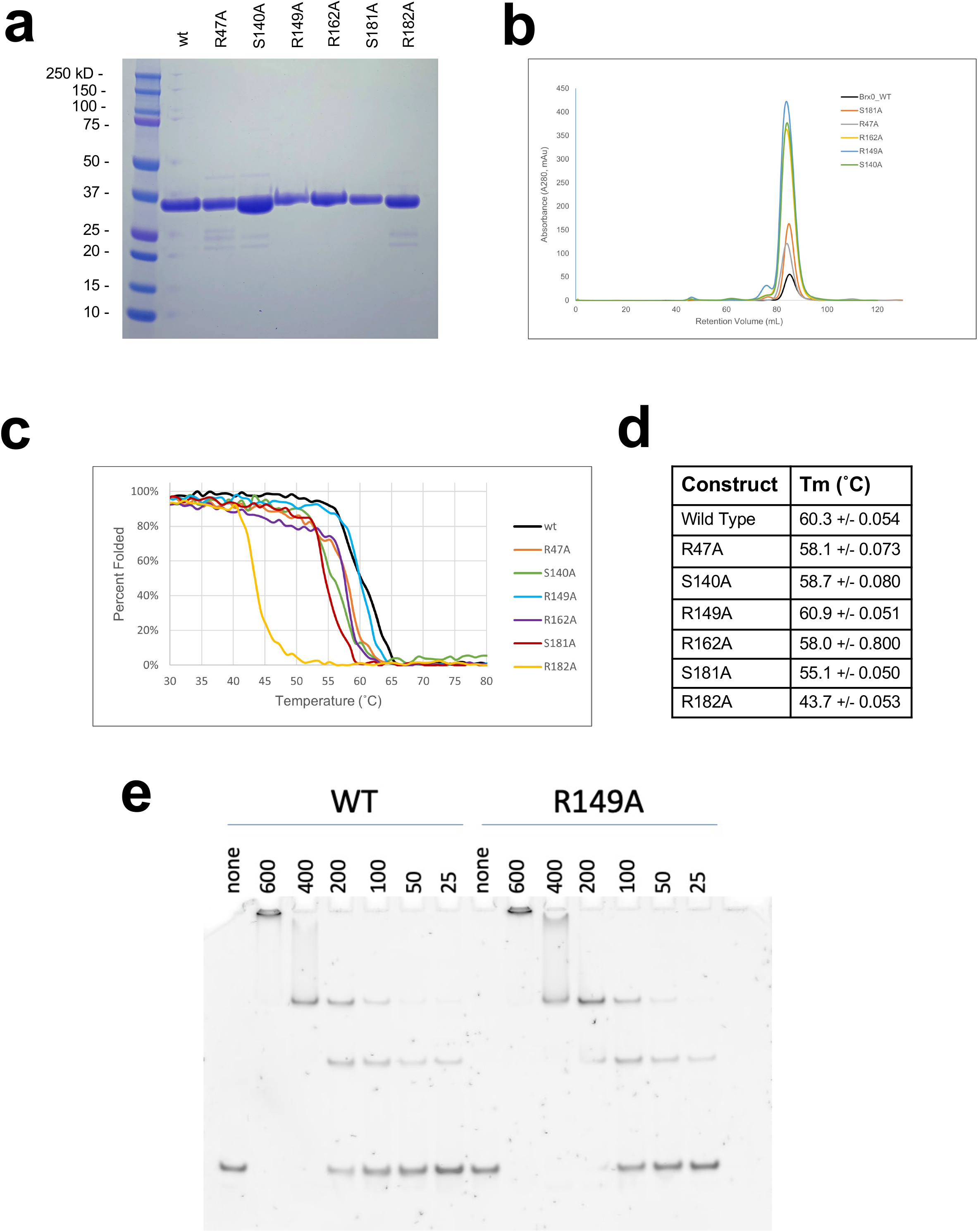
Purification and characterization of BrxR point mutants. A series of six point mutations (R47A, which is located in the protein’s DNA binding domain, and five additional constructs located in the WYL domain) were purified (***panel a***) and demonstrated to all elute at volumes corresponding to dimers on a size exclusion column (***panel b***). Subsequent determination of their thermal denaturation temperatures and unfolding behavior using CD spectroscopy demonstrated that one point mutant (R182A) was significantly destabilized (***panels c-d***). One point mutant in the WYL domain (R149A) was analyzed for DNA target binding and shown to interact with that site in a manner similar to the wild-type protein (***panel e***). The first three lanes in panel a are also shown in Figure 8a; in both cases they illustrate the outcome from the same experiment (purification of WT and R47A BrxR).

**Supplementary Figure S10.**
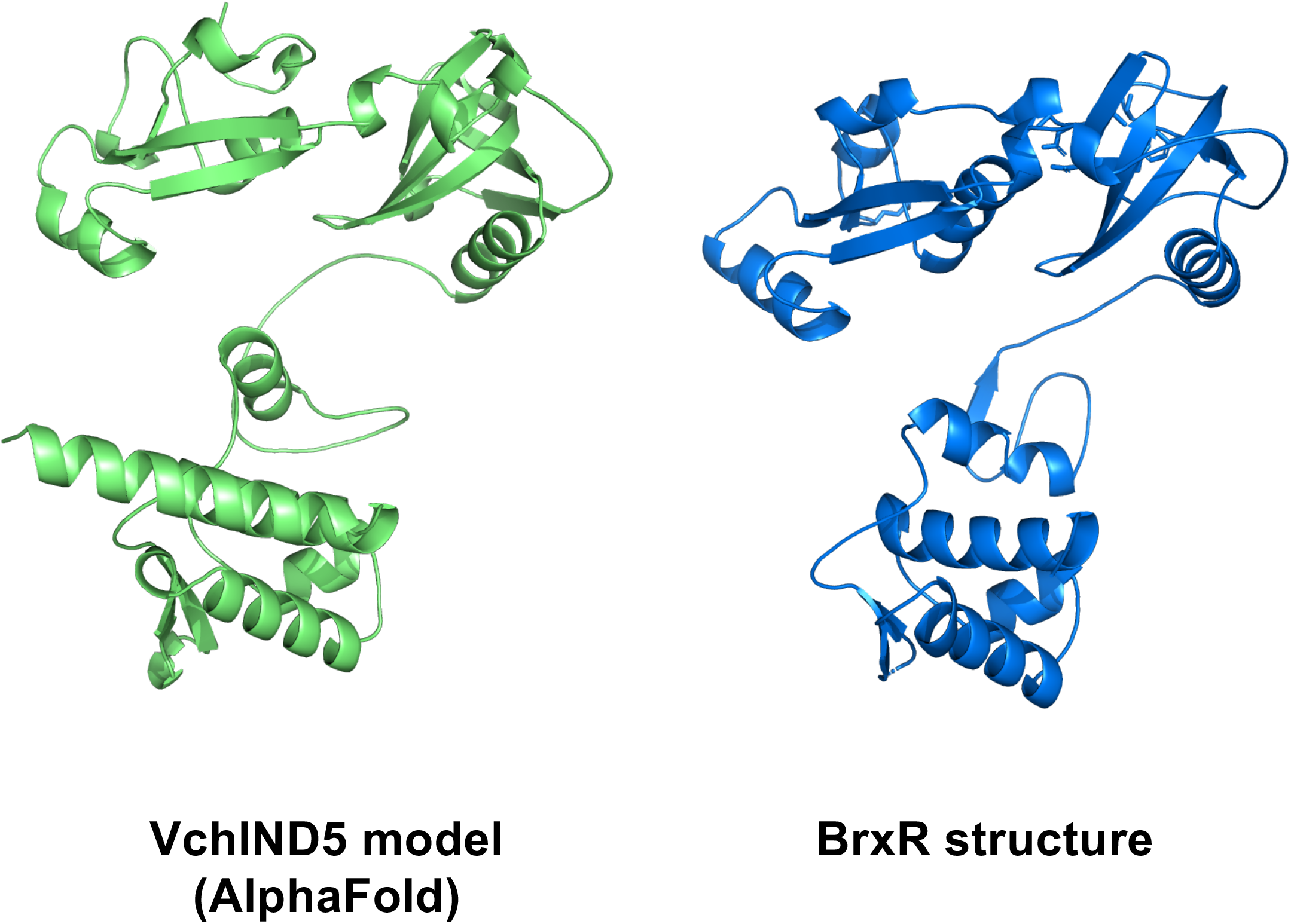
WYL proteins upstream of defense islands in *V. cholera* are BrxR homologs. VchInd5, which was annotated by Legault et al as a “WYL” protein, shares ∼25% amino acid sequence identify with BrxR. The predicted Alpha-fold structure of VchInd5 (left) closely matches the experimentally determined structure of BrxR, demonstrating that the V. cholera WYL proteins are BrxR homologs.

---

**Supplementary Table S1.**
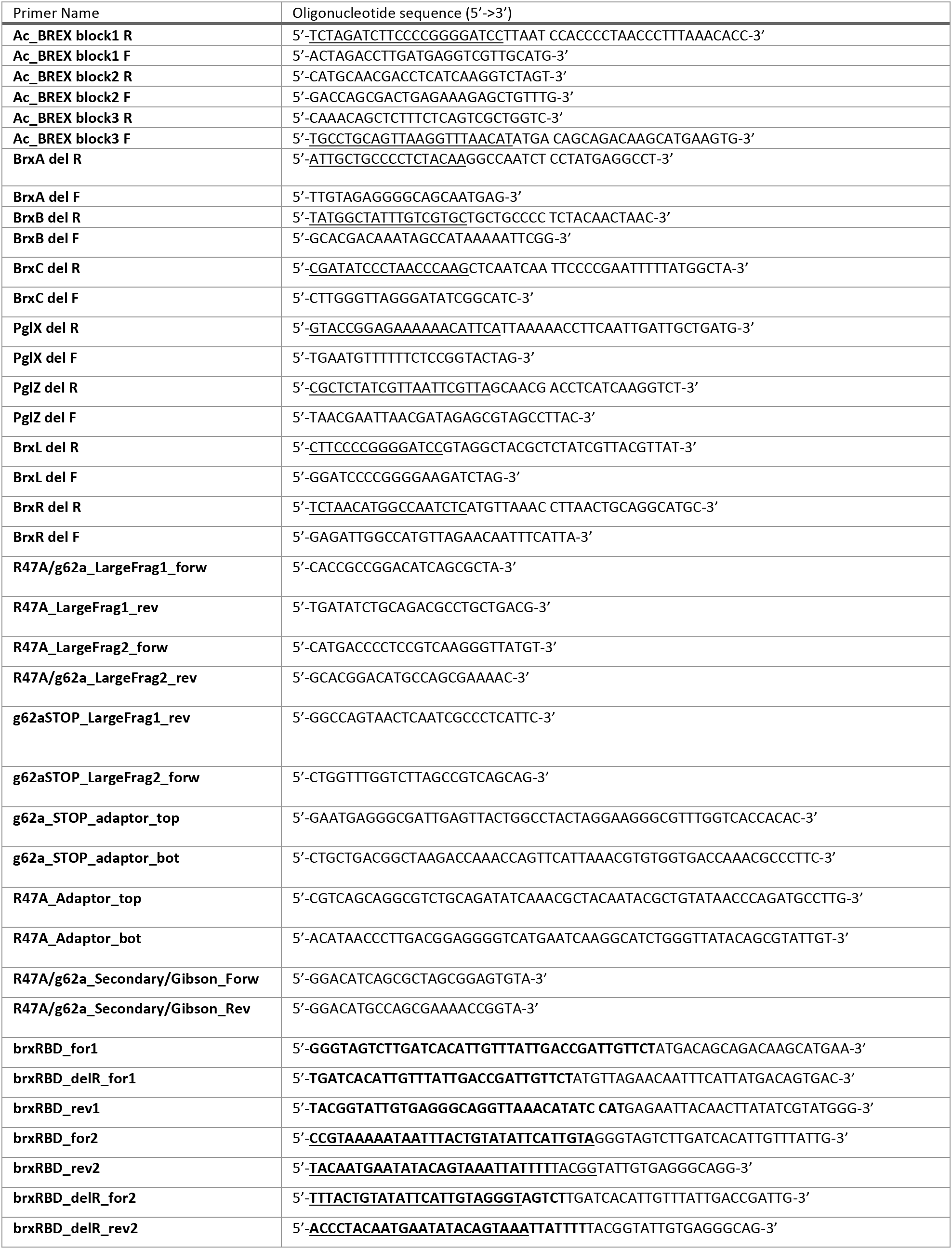
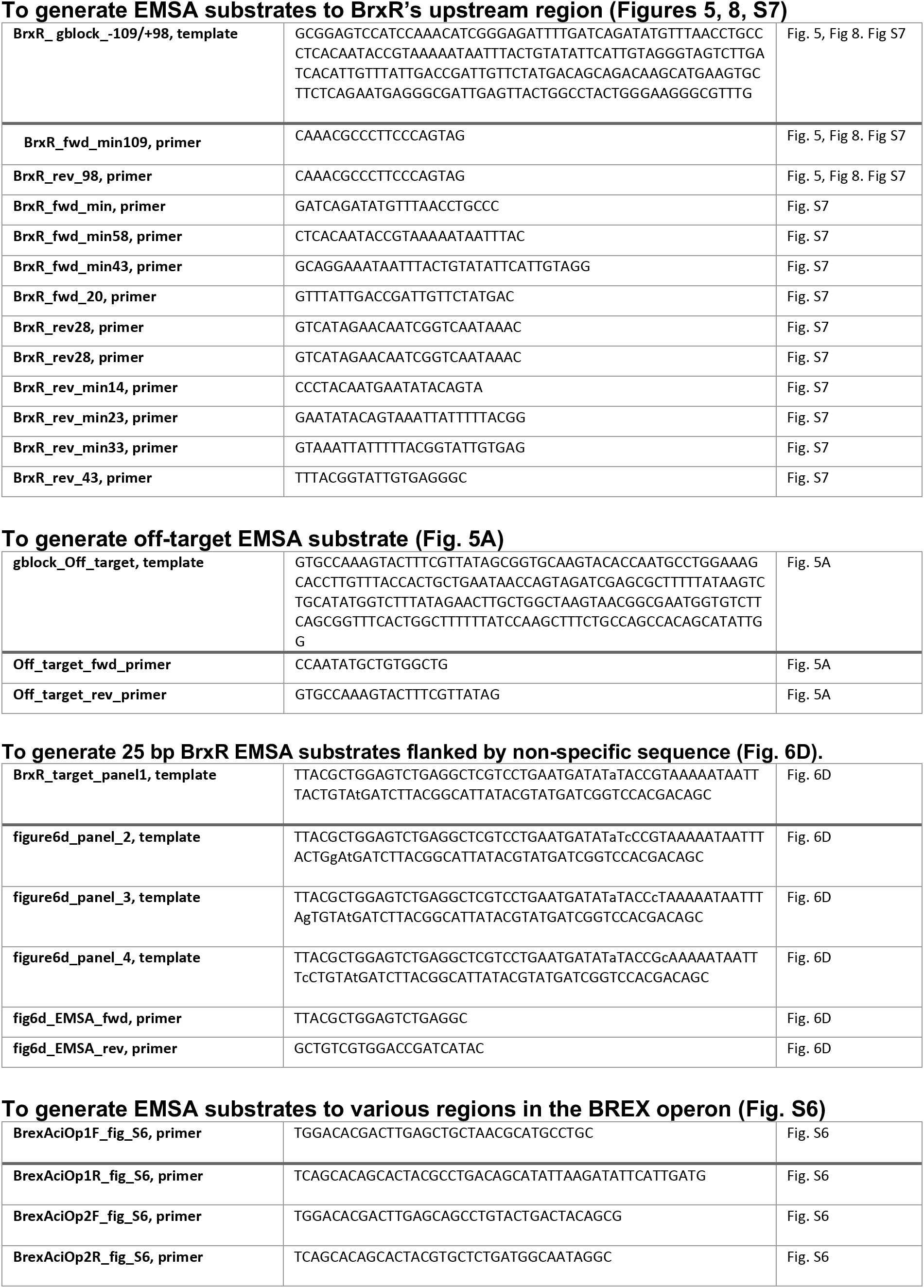
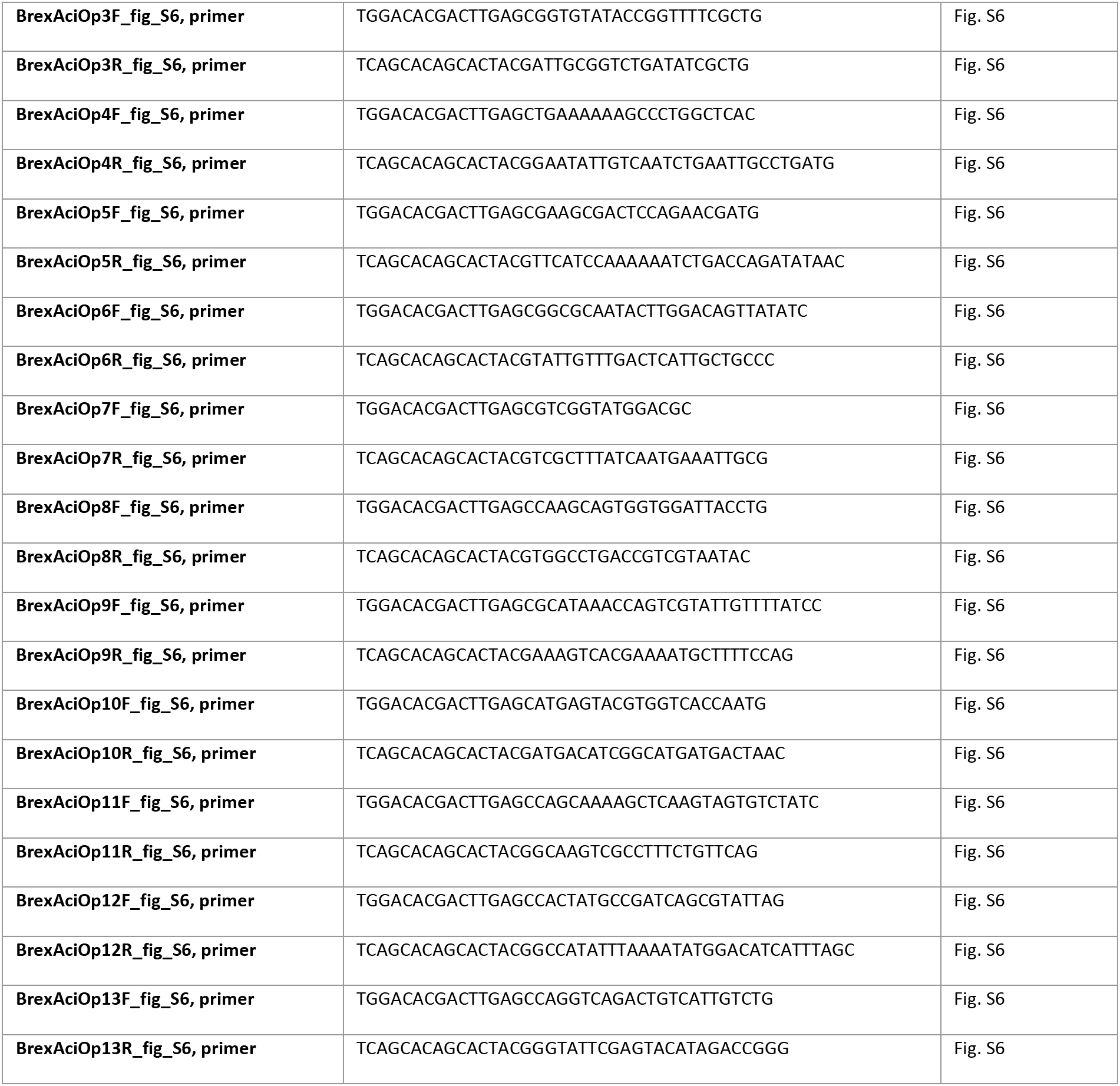
Primers for Molecular Biology and Subcloning and DNA constructs for binding assays.

